# Most of transcriptional alterations in glioma result from DNA-methylation independent mechanisms

**DOI:** 10.1101/516997

**Authors:** Franck Court, Elisa Le Boiteux, Anne Fogli, Mélanie Müller-Barthélémy, Catherine Vaurs-Barrière, Emmanuel Chautard, Bruno Pereira, Julian Biau, Jean-Louis Kemeny, Toufic Khalil, Lucie Karayan-Tapon, Pierre Verrelle, Philippe Arnaud

## Abstract

**Background:** Aberrant DNA methylation is a cancer cell feature that is commonly associated with transcriptional alterations. However, the primary role of this defect in the genome-wide cancer-associated gene deregulation is not clear. Here, we evaluated the relative contribution of DNA methylation-dependent and -independent mechanisms to transcriptional alterations at CpG-island/promoter-associated genes in samples of adult glioma, a widespread type of brain tumor.

**Results:** Extensive molecular analyses of glioma samples with wild type IDH (IDHwt) and mutated IDH (IDHmut) found DNA hypermethylation only in a minority of genes showing loss or gain of expression. Specifically, in IDHwt samples, more than 75% of aberrantly repressed genes did not display DNA methylation defects at their CpG-island promoter. Conversely, altered H3K27me3 was the predominant molecular defect at deregulated genes. Moreover, the presence of a bivalent chromatin signature at CpG-island promoters in stem cells might not only predispose to hypermethylation, as widely documented, but more generally to all types of transcriptional alterations in transformed cells. In addition, the gene expression strength in healthy brain cells influences the choice between DNA methylation- and H3K27me3-associated silencing in glioma. Our findings support a model whereby the altered control of H3K27me3 dynamics, more specifically defects in the interplay between polycomb protein complexes and the brain-specific transcriptional machinery, is the main cause of transcriptional alteration in glioma cells.

**Conclusions:** Our study provides a comprehensive and updated description of the epigenetic deregulations in glioma that could be useful for the design of drugs to target cancer-related epigenetic defects.

## Background

Cancer is a complex disease that results from the disruption of key pathways, including those regulating cell survival and division. Besides genetic lesions, epigenetic alterations also contribute to tumorigenesis mainly by leading to abnormal gene expression (1).

Together with genome-wide DNA hypomethylation, DNA hypermethylation of CpG islands (CGIs) is a well-defined feature of cancer cells, and is believed to be the main cause of aberrant gene repression (2). CGIs are key regulatory genomic regions of few hundred base pairs in size characterized by high frequency of CpG dinucleotides. In humans, about 70% of promoters are associated with CGIs that generally remain unmethylated during somatic development, regardless of the gene expression status (3). Conversely, it has been shown that in tumors, DNA hypermethylation of their CGI/promoter leads to aberrant silencing of some tumor suppressor genes, such as *BRCA1* (*4), RB1 (5)* and *MLH1 (6)*. However, the primary role of this defect in widespread cancer-associated genes silencing, and more broadly in cancer biology, is still questioned. Indeed, an increasing number of studies showed that in tumors, DNA hypermethylation affects primarily CGI/promoters that control genes already repressed in the matched normal tissue (7–11). Moreover, in some tumor types, such as glioma or breast cancer, patients with a CpG island methylator phenotype (CIMP), a signature identified in various human malignancies and defined by the concomitant hypermethylation of multiple CGIs (12), have a better clinical prognosis compared with patients without CIMP (13–14).

Therefore, other DNA methylation-independent epigenetic alterations at CGI/promoters might contribute to the genome-wide pattern of aberrant gene repression observed in cancer cells. During normal development, promoters/CGIs are dynamically marked by the permissive H3K4me3 and/or the repressive H3K27me3 histone marks (when in combination, they constitute the so-called bivalent chromatin signature) (15). Alterations in the control of these chromatin signatures also could lead to gene silencing (11). This hypothesis is further supported by the observation that methyltransferases and demethylases that regulate H3K27me3 and H3K4me3 deposition, such as EZH2, the MLL family members and UTX, are translocated or mutated and/or their expression is altered in many malignancies (16,17). Such defects had been documented in a handful of studies. In detail, analyses in established prostate (*7, 18*) and urothelial (19) cancer cell lines and in colorectal tumor samples (20) highlighted that gene silencing can be mediated just by H3K27me3.

Chromatin-based alterations could also lead to gain of gene expression in tumors. Hahn et al. showed that genes associated with GCI/promoters displaying a bivalent chromatin signature in normal colon can be ectopically expressed in the matched tumor samples following the loss of H3K27me3 (20). Moreover, Bert et al. (21) identified in prostate cancer cell lines some genomic domains characterized by altered chromatin signatures associated with aberrant gene expression.

Therefore, it is clear that various epigenetic alterations at promoters/CGIs could contribute to the abnormal gene expression pattern of cancer cells. Because of the absence of dedicated integrative studies, many questions remain concerning the bases and extent of these alterations as well as their relative contribution to aberrant loss/gain of gene expression in cancer cells.

Here, we used glioma as a model to investigate the molecular bases of the transcriptional alteration of CGI/promoter-associated genes in cancer. Glioma is derived from glial cells, and is one the most widespread brain tumor types. In 2007, the World Health Organization (WHO) classified gliomas in four grades (I-IV) according to their histology. Malignant anaplastic astrocytoma (a subset of the WHO grade III gliomas) and glioblastoma multiforme (GBM; WHO grade IV) account for about half of all gliomas, and are the most deadly and aggressive forms. The median survival time after diagnosis of patients with GBM does not exceed 18 months despite the aggressive treatments. At the molecular level, aggressive gliomas are characterized by expression of wild type isocitrate dehydrogenase (IDHwt) genes (*IDH1* and *IDH2*), while gliomas with better prognosis express mutated *IDH* (IDHmut) (22). Consequently, the recently released 2016 WHO classification of diffuse gliomas (23), which we used in this study, is primarily based on the *IDH* mutation status (IDHmut *vs* IDHwt). Strikingly, *IDH1* mutation results in tumors with a CIMP (24), and with a better clinical prognosis compared with CIMP-negative tumors (13, 22). Other epigenetic regulators also could be involved in glioma development/progression, for instance the polycomb repressors EZH2 and BMI1 (25–27) and MLL family members that are mutated in a subset of GBM (28, 29).

The present integrative molecular analysis of well-characterized IDHwt and IDHmut glioma samples highlights that alteration of the H3K27me3 signature, rather than of DNA methylation, is the predominant molecular defect associated with aberrantly repressed and expressed genes.

## Results

### CGI methylation poorly contributes to transcriptional alterations in glioma

For this study, we used 70 clinically well-characterized primary glioma samples from Clermont-Ferrand Hospital, France (the patients’ demographic and main molecular and clinical features are provided in Additional file 1: Table S1). We classified samples according to their *IDH1* status (*n* = 55 IDHwt, and *n* = 15 IDHmut). This first level of classification, upstream 1p/19q codeletion status according the 2016 WHO classification (23), indeed clearly discriminated two tumor classes relative to aggressiveness, with a significant survival advantage for patients with IDHmut compared with IDHwt tumors (HR=0.32, 95% CI [0.14-0.71], p=0.005) (Additional file 1: Table S1 and Figure S1a). Genome-wide analysis of DNA methylation at CGI areas, using the Infinium HumanMethylation450 (HM450K) BeadChip Arrays, showed that DNA methylation defects were more widespread in IDHmut samples, constituting a Glioma CIMP (G-CIMP) subclass (Additional file 1: Figure S1b). In agreement with the literature (13, 23, 24), in our cohort, aggressive gliomas were characterized by IDHwt and absence of G-CIMP, whereas IDHmut gliomas showed a G-CIMP profile and had a better prognosis.

To more precisely define the CGI/promoter alterations in our glioma samples, we analyzed the DNA methylation profiles of 14714 genes with a single CGI-rich promoter that could be assessed using the HM450K array. Most of these genes (76%) were protein-coding genes, and the others were antisense transcripts (10.6%), long intergenic non-coding RNAs (lincRNA; 6.5%), and pseudogenes (6.8%) (Fig. 1a). As some CGI/promoters can control more than one gene, these 14714 genes are associated with 11795 CGI/promoters. About 90% of these CGI/promoters were unmethylated in non-tumor control brain samples (mean β-value <0.2) and most of them remained unmethylated also in glioma samples. Compared with control samples, only 11.6% of GCI/promoters (n=1369; associated with 1623 genes) were aberrantly methylated in IDHwt samples, and 22.8% (n=2692; 3198 genes) in IDHmut samples, contributing to the CIMP-positive status of these samples. Conversely, though much more limited CGI/promoter hypomethylation was more common in IDHwt (n=198 CGI/promoters associated with 235 genes; 1.7%) than in IDHmut (n=14 CGI/promoters associated with 22 genes; 0.12%) samples (Fig. 1b; Additional file 1: Figure S1c-d).

**Figure 1:**
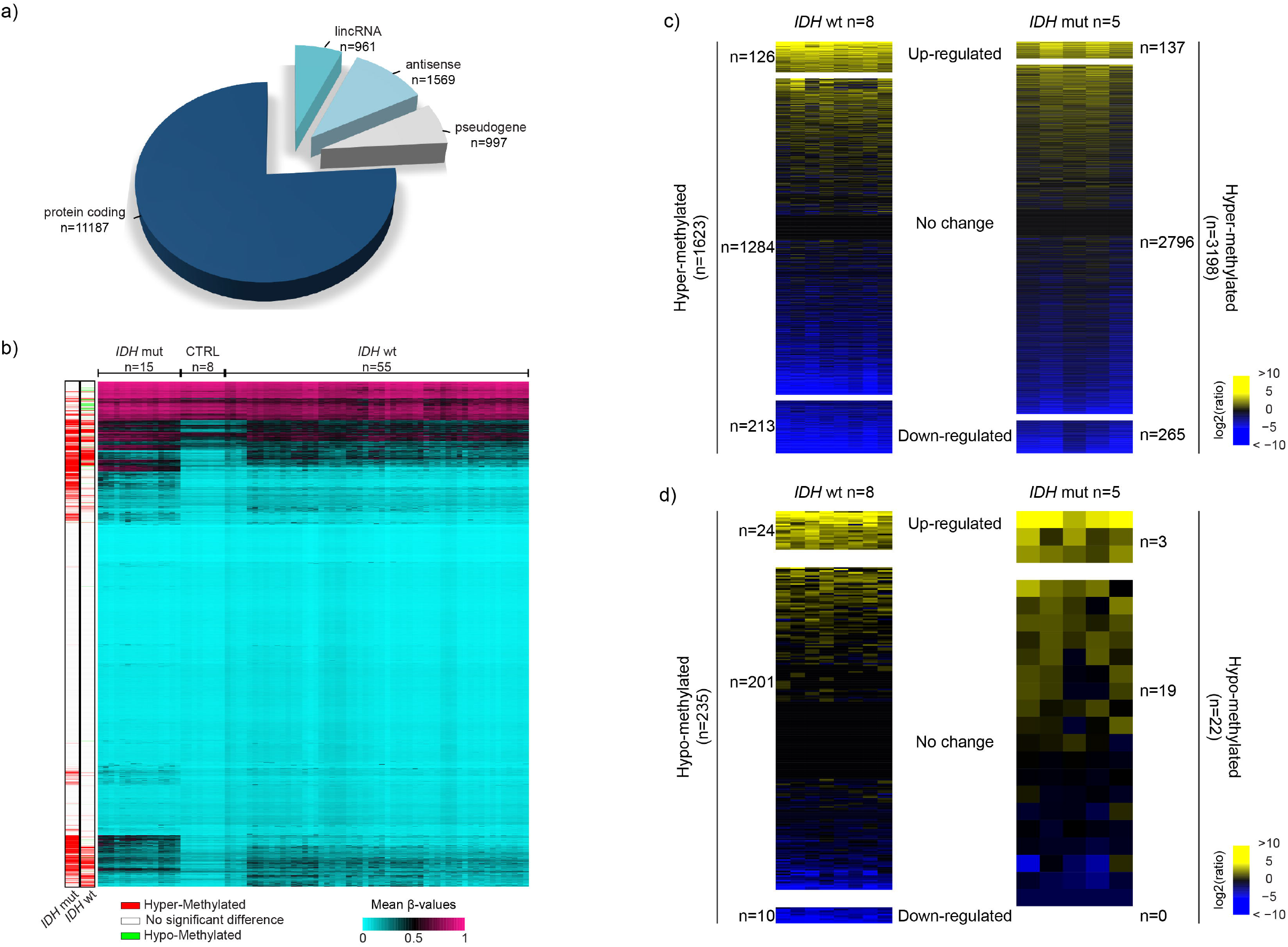
Aberrant methylation at CGI/promoters is not the main contributor to transcriptional alteration in glioma. **a)** Classification of the 14714 genes analyzed in this study. **b)** DNA methylation level (mean β-values) of the 11795 CGI/promoters (row) analyzed in IDHmut and IDHwt glioma and control (normal brain tissue) samples (columns). Left columns show their hyper- or hypo-methylation status in IDHmut and IDHwt glioma samples compared with controls. **c-d)** Differential expression status of genes associated with hyper- **(c)** or hypomethylated **(d)** CGI/promoters in IDHwt (left panel) and IDHmut (right panel) glioma samples compared with controls.

To evaluate the consequence of these DNA methylation defects, we analyzed the RNA-seq data we obtained from 8 IDHwt and 5 IDHmut glioma samples. About 80% of genes with aberrantly methylated CGI/promoters in the IDHwt and 87.5% in the IDHmut group did not show significant transcriptional change (|Log2 Fc|>2; p<0.05) (Fig 1c,d). Therefore, and despite the marked difference in DNA methylation profiles, the number of affected genes was similar between glioma subtypes. Among the genes with aberrantly methylated CGI/promoters, 223 and 265 were downregulated (82 in common), and 150 and 140 genes were upregulated (44 in common) in IDHwt and IDHmut glioma samples, respectively, compared with controls (Fig 1c,d). These findings indicate that aberrant CGI/promoter methylation minimally affects gene transcription in glioma. However, the deregulated genes included some putative tumor suppressors, such as *RRP22* and *HTATIP2* (30, 31), and some putative oncogenes, such as *HOXD9* and *CXCL1*, the overexpression of which was, counterintuitively, associated with methylation gain (Additional file 2: Table S2).

### Gene transcription alterations are more widespread in IDHwt glioma samples

Detailed analysis of the transcriptional landscape of tumor samples showed that transcriptional alterations were more widespread in IDHwt than in IDHmut glioma samples (1670 and 1024 deregulated genes, respectively; FDR<0.05; |Log2Fc|>2) (Fig. 2a), particularly for CGI/promoter-associated genes (Additional file 1: Figure S2a).

**Figure 2:**
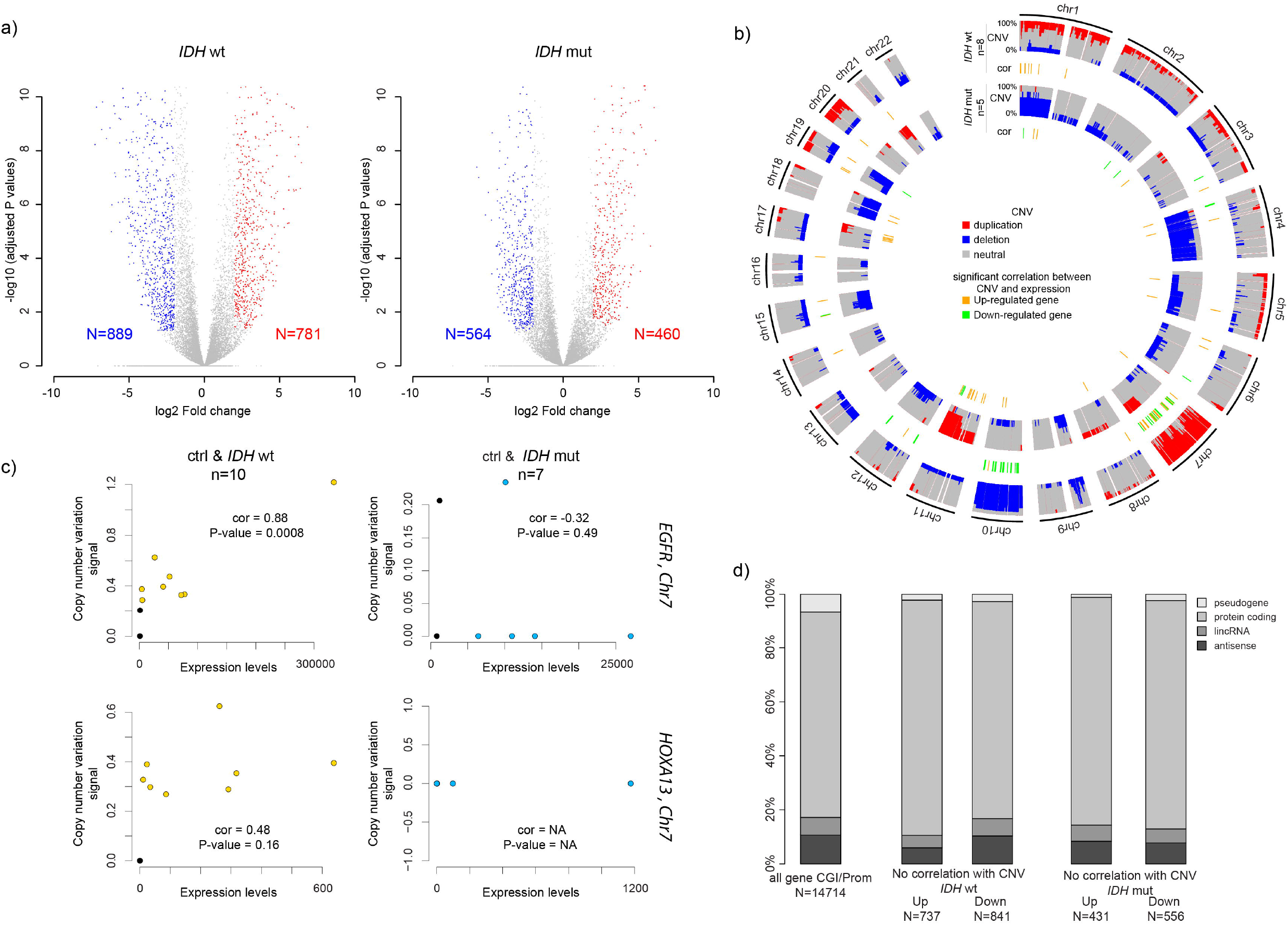
Extent of transcriptional alterations in IDHwt and IDHmut glioma samples. **a)** Volcano plot analysis of differential gene expression in IDHwt (left) or IDHmut (right) glioma samples. Blue and red dots represent genes that were significantly down- or up-regulated, respectively, compared with healthy controls (n=14714 genes analyzed). **b)** Circular karyotype showing the duplication (in red) and deletion (in blue) frequencies at the 14714 analyzed genes in IDHwt (outer circles) and IDHmut (inner circles) samples. Genes showing a significant correlation between CNV and expression are symbolized by an orange (up-regulated) or green (down-regulated) line. **c)** Correlation analysis between CNV and expression levels for the *EGFR* and *HOXA13* genes in IDHwt (yellow dots, left panels) and IDHmut (blue dot, right panels) glioma samples. Black dots indicate value in healthy controls. *EGFR* overexpression correlated with increased copy number in IDHwt glioma samples. **d)** Classification of the genes with expression alterations that did not correlate with CNV.

Copy number variation (CNV) analyses in the same samples showed that, as previously reported, chromosome 7 gain and chromosome 10 loss characterized IDHwt samples (23), while the 1p/19q codeletion was mainly present in IDHmut samples (Fig 2b). By integrating these data with the gene expression profiles, we identified 92 genes in IDHwt and 37 genes in IDHmut samples, respectively, in which expression alteration correlated with CNV (p<0.05) (Fig 2b, and Additional file 1: Figure S2b-c, Additional file 3: Table S3). For instance, upregulation of epidermal growth factor receptor (*EGFR*) and the histone methyltransferase *EZH2* (both located on chromosome 7) correlated with increased copy number in IDHwt samples (Fig 2c; Additional file 1: Figure S2c). Conversely, overexpression of *HOXA13*, also located on chromosome 7, did not correlate with CNV.

Altogether, affected genes without CNV (mostly protein-coding genes) represented about 11% of all CGI/promoter-associated genes in IDHwt samples (841 downregulated and 737 upregulated genes), and 6.7% in IDHmut samples (556 downregulated and 431 upregulated) (Fig 2d, Additional file 2: Table S2).

### Most transcriptional alterations are DNA methylation-independent

To understand the bases of transcriptional alteration in IDHwt glioma samples, we used paired RNA-seq and HM450K data from 8 IDHwt samples to concomitantly determine the DNA methylation and transcriptional changes in the 1578 affected genes. In these IDHwt samples, we could identify four main transcriptional defect types (Fig. 3a): gain or loss of gene expression associated with CGI/promoter hypermethylation (defect 1 and 3, respectively), and gain or loss of gene expression without changes in CGI/promoter methylation status (*i.e.*, the CGI/ promoter remained unmethylated) (defect 2 and 4, respectively). More than 93% of aberrantly repressed genes did not display any significant DNA methylation alteration at their CGI/promoter (defect 4). About 47% of the affected genes showed gain of expression that was associated with DNA hypermethylation at their CGI/promoter in 6% of them (defect 1).

**Figure 3:**
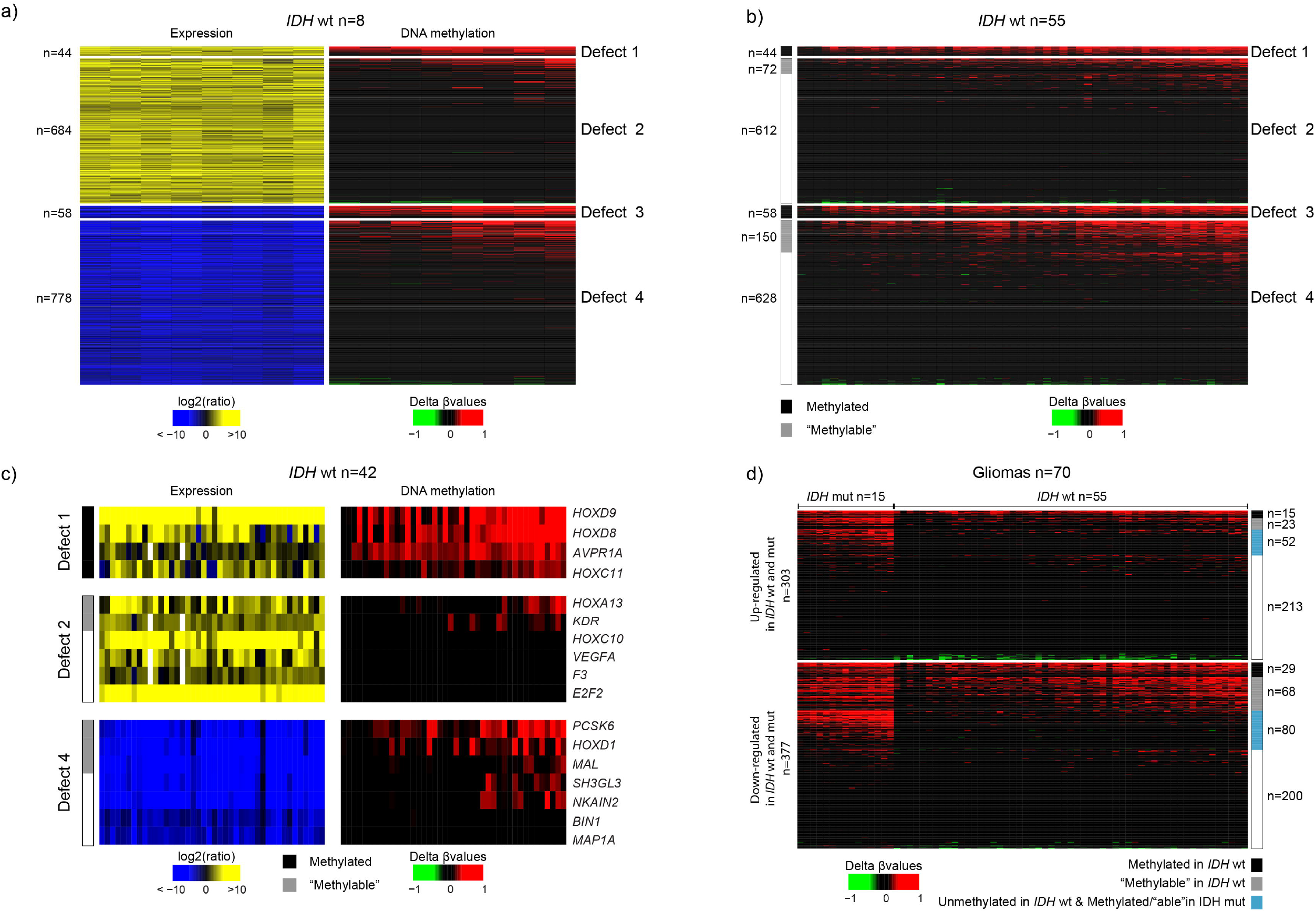
Four expression defect classes. **a)** Integrative analysis of differential gene expression and methylation in 8 IDHwt glioma samples identified four main defect classes. **b)** Differential DNA methylation analysis in all IDHwt glioma samples (n=55) ra controls (n=8) (delta of the mean ß-value). Glioma samples were grouped in the four classes of expression defects defined in a). The methylated and methylable status of genes is indicated in the left column. **c)** Integrative analysis of differential expression and methylation at selected genes with defect 2 (upper panel), 3 (middle panel) and 4 (lower panel) in 42 IDHwt glioma samples compared with controls (n=8). **d)** Differential methylation pattern, vs healthy controls, of genes affected both in IDHwt and IDHmut glioma samples. Genes identified as methylated and methylable in IDHwt glioma samples (b) are shown in black and grey, respectively, on the right column. Genes unmethylated in IDHwt glioma samples and methylated/methylable in IDHmut glioma samples are shown in blue.

To test the robustness of this classification, we first analyzed the HM450K data of all 55 IDHwt glioma samples of our cohort and confirmed that DNA hypermethylation was associated with genes classified in defects 1 and 3, and hypomethylation with those classified in defects 2 and 4. The only exception was a subset of “methylable” genes that could gain methylation in some samples (Fig. 3b) and thus, displayed an overall significant gain of DNA methylation. Next, we concomitantly assessed, in 42 IDHwt glioma samples, the DNA methylation and expression, by RT-qPCR, of randomly selected genes from defect 1,2 and 4 group, respectively. The seven genes from defect 4 group were aberrantly repressed in all analyzed samples and their CGI/promoter mostly unmethylated (Fig. 3c). For instance, the candidate tumor suppressor gene *BIN1* was unmethylated in all analyzed samples. *PCSK6* and *HOXD1* provided examples of “Methylable” genes. They were methylated in a subset of samples, but their expression was repressed in all of them. The six genes from defect 2 group were all overexpressed and most of them, including the tumor progression-associated *VEGFA* and *E2F2* genes, tended to be unmethylated in all analyzed samples. Defect 2 group also included some “methylable” genes (see Fig. 3b), such as *KDR* (the tyrosine kinase receptor for VEGFA) that was overexpressed in all samples, whereas its CGI/promoter was methylated only in a subset of gliomas. Genes with defect 1 were methylated and ectopically expressed in all IDHwt glioma samples.

To further characterize these defects, we studied the methylation pattern of the 681 genes that were transcriptionally affected in both IDHwt and IDHmut glioma samples (377 downregulated and 303 upregulated in both groups, and 1 downregulated in IDHwt and upregulated in IDHmut). Most of these genes displayed the same methylation pattern in IDHwt and IDHmut samples. However, “methylable” genes and a subset of unmethylated genes in IDHwt gliomas (symbolized by a blue column in Fig. 3d) tended to be methylated in IDHmut samples, suggesting that different molecular pathways can lead to the same aberrant gene expression pattern.

Altogether, this integrative analysis identified four classes of transcriptional defects at CGI/promoter genes in IDHwt glioma: aberrant loss and gain of gene expression without DNA methylation defect (most genes), and gene expression defects associated with constitutive or optional (*i.e.*, methylable genes) aberrant methylation. Consequently, we reclassified the “methylable” genes that were initially included in the defect 2 and 4 groups into the defect 1 and 3 groups, respectively, for the subsequent analyses. In summary, among the 1578 CGI/promoter-associated genes affected in IDHwt glioma samples, most belonged to the defect 4 group (n=628), followed by defect 2 (n=612), defect 3 (n=208), and finally defect 1 group (n=116). The classification for each gene is provided in Additional file 2: Table S2.

### The four defect classes include mainly genes with a chromatin bivalent signature in stem cells

Gene ontology analysis (Fig. 4a, Additional file 1: Figure S3) showed that defect 3 and 4 groups (gene repression associated or not with DNA hypermethylation) were enriched in genes involved in transmembrane ion transport, specifically in synapsis and neurons (defect 3: repression without DNA hypermethylation). Conversely, defect 2 group (gene overexpression) were enriched in genes implicated in general processes, such as cell cycle, cell division and chromosome segregation. Moreover, we found that the defect 1 group (overexpression with methylation) was highly enriched in gene encoding homeodomain proteins, including the HOX family, and implicated in embryonic development.

**Figure 4:**
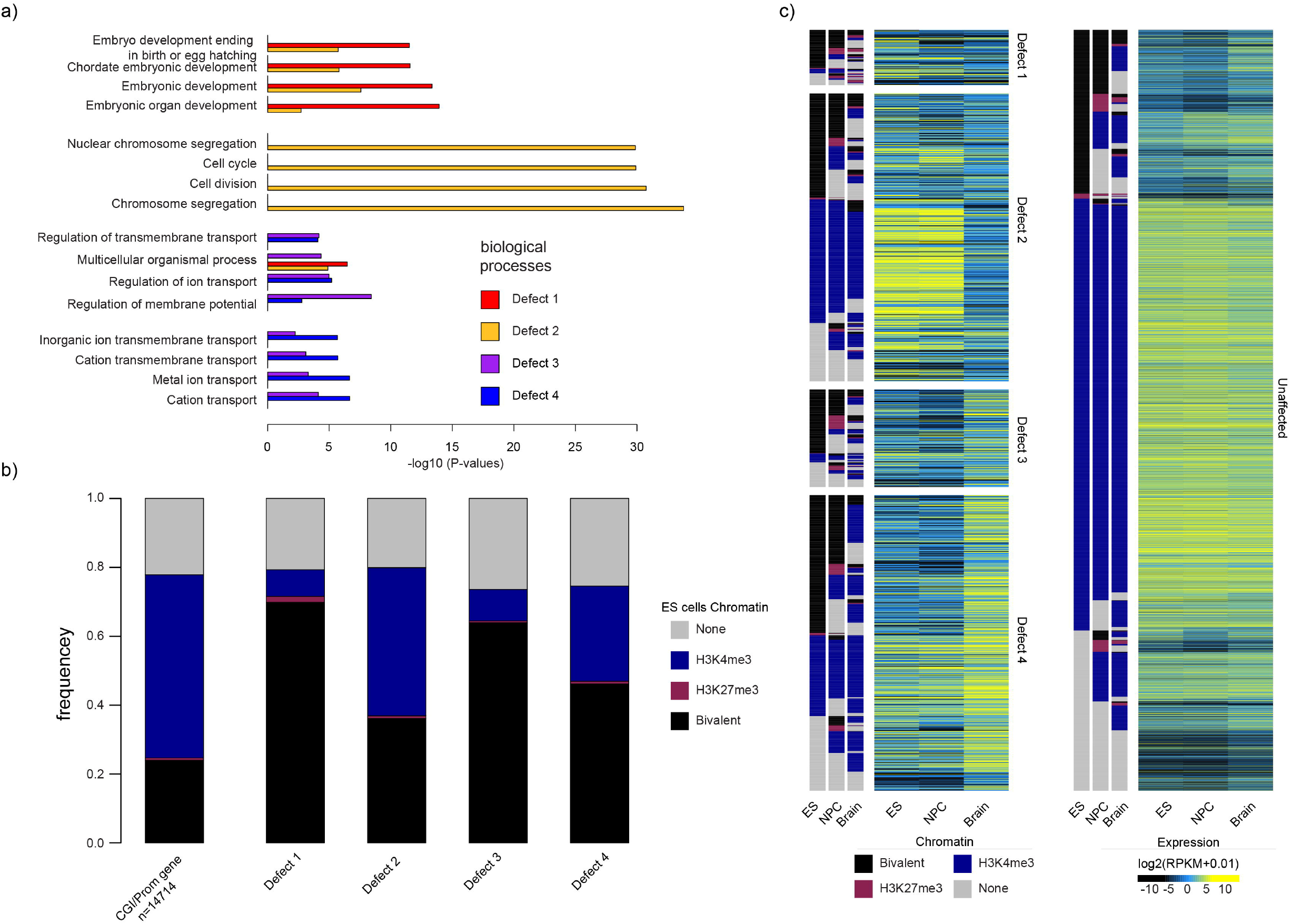
Genes with bivalent chromatin signature in ES cells are more prone to be deregulated in IDHwt glioma. **a)** Gene ontology terms (biological processes) enriched in genes with defect 1 to 4. For each category, the four highest terms are shown. **b)** Distribution of genes with defect 1 to 4 according to their chromatin signature in human ES cells (none: gray; bivalent: black; H3K4me3-only: blue; H3K27me3-only: red). As reference, the distribution of the 14714 genes analyzed in this study according to their chromatin signatures in human ES cells is shown in the left panel. **c)** Expression level and chromatin signatures of genes with defect 1 to 4 in human ES cells, neural progenitor cells (NPC) and healthy brain. For comparison, the same analysis is provided on the right panel for genes without expression defect (unaffected) in IDHwt glioma samples.

We next evaluated the GCI/promoter chromatin signature of genes in these four categories, in human embryonic stem (ES), in neural progenitor cells (NPC) and brain samples). In ES cells and NPC, genes in the defect 1 and 3 groups showed a bivalent chromatin signature. This is in agreement with previous findings showing that genes with a bivalent chromatin signature in stem cells are more likely to gain aberrant methylation in cancer cells (11, 32, 33). Similarly (but more unexpectedly), about 36% and 48% of genes in the defect 2 and 4 groups, respectively, showed a bivalent chromatin signature in ES cells (compared with 24% of all studied genes) (Fig. 4b, Additional file 1: Figure S3). In agreement with the resolution of the bivalent signature during development/cell differentiation, the chromatin signature tended to change towards an exclusive H3K4me3 signature, but also to a “none” signature (i.e: depleted for both H3K4me3 and H3K27me3) in brain samples (Fig. 4c). In comparison, most of the transcriptionally unaffected genes maintained their H3K4me3-only signature from ES cell to brain samples. Accordingly, the transcriptionally affected genes displayed a dynamic expression pattern from ES cells to NPC and brain. Interestingly, this was true also for the subset of genes in the defect 2 and 4 groups that maintained an exclusive H3K4me3 signature in ES cells, NPC and brain, and nonetheless showed a loss and gain of expression in brain, respectively (Figure 4 c).

Altogether, these findings suggest that genes with a bivalent chromatin signature in ES cells and/or with a dynamic expression pattern during neural differentiation are more prone to be deregulated in IDHwt glioma.

### CGI/promoter hypermethylation is associated with gain of expression

To determine how gain of methylation and expression could co-exist in genes with defect 1, we evaluated their CGI/promoter chromatin signature using publicly available ChIP-seq data. Compared with healthy brain (control), H3K27me3 level was strongly decreased in IDHwt GBM samples, while H3K4me3 level was increased in about 65% of glioma samples and totally depleted in the others (Fig. 5a).

**Figure 5:**
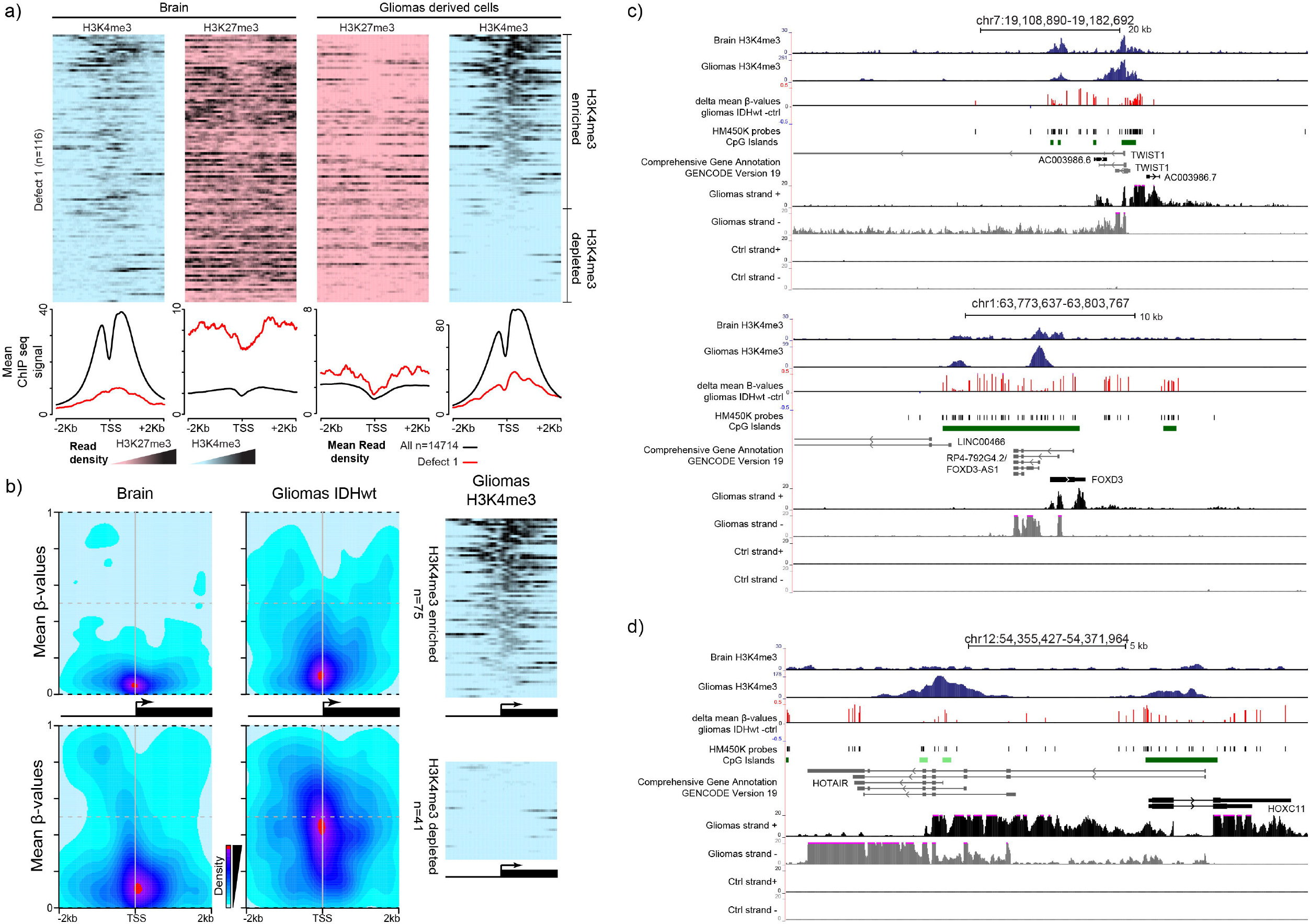
Expression from genes with methylated CGI/promoter. **a)** Data-mining derived ChIP-seq read density data for H3K27me3 (purple) and H3K4me3 (blue) in genes with defect 1 in a ± 2Kbp window centered on their TSS, in healthy brain (left panel) and IDHwt-derived cell lines (right panel). The mean ChIP-seq signal values are shown on the lower panels for genes with defect 1 (red line) and for the 14714 analyzed genes (black line) that were used as normalized reference. **b)** Heatmap showing CpG sites density and their mean methylation level in a ± 2Kbp window centered on the TSS of genes with defect 1 and enriched (upper panel) or depleted (lower panel) for H3K4me3 in IDHwt glioma samples, compared with healthy controls. The ChIP-seq read density obtained in IDHwt-derived cell lines is shown on the right panels. **c)** Genome browser view at the *TWIST1* and *FOXD3* loci to show H3K4me3 enrichment, differential DNA methylation, and the oriented RNA-seq signal. These two loci are representative of genes that initiate from an H3K4me3-marked TSS embedded in a methylated CGI/promoter in IDHwt samples. **d)** *HOX-C11* is representative of genes in which expression initiates from an alternative TSS in IDHwt glioma samples.

Analysis of the HM450K data on the localization of hypermethylated sites relative to the transcription start site (TSS) showed that at H3K4me3-enriched genes, the gain of DNA methylation occurred at the border of the CGI/promoter, while the TSS area remained unmethylated (Fig. 5b). Analysis of individual loci in glioma samples using also strand oriented RNA-seq data also suggested that transcription initiates from H3K4me3-marked TSS embedded in methylated CGIs (Fig. 5c, and Additional file 1: Figure S4a). This was observed for several genes that promote gliomagenesis, including *TWIST1* (34), *CTHRC1* (35) and *FOXD3-AS1* (36). The H3K4me3 and DNA methylation signals were mutually exclusive (Fig. 5c and Additional file 1: Figure S4a), in agreement with the documented antagonism of these two marks (37).

In H3K4me3-depleted genes, DNA methylation was spread along the entire CGI/promoter, including the TSS (Fig. 5b), suggesting that transcription from these genes could arise from an alternative TSS. Analysis of RNA-seq data supported this hypothesis because genes, such as *HOXC11* and *NRF2*, showed transcription signal from H3K4me3-enriched regions located away from the documented TSS (Fig 5d and Additional file 1: Figure S4b). However, for few genes, such as *HEYL* and *C15orf48* (Additional file 1: Figure S4c), transcription apparently initiated from a methylated CGI through an unknown mechanism.

Altogether, these approaches support the hypothesis that in genes with defect 1, transcription could be allowed by the absence of DNA methylation at the TSS, or the use of alternative TSS.

### E2F- and HOX-target genes are frequently overexpressed in glioma

The publicly available ChIP-seq data indicated that genes with defect 2 were enriched in H3K4me3 and depleted in H3K27me3 in glioma samples compared with healthy brain (Fig. 6a). We observed this H3K4me3 gain also at the subset of genes that were constitutively marked by the H3K4me3-only signature in ES cells, NPC and brain (framed in Fig. 6a). To understand the basis of their overexpression in glioma, we asked whether specific motifs were enriched at their CGI/promoters. We found that overall, genes with defect 2 were putative targets of transcription factors associated with cell cycle pathways, including the Krüppel-like factors (KLF)/specificity protein (SP) and E2F families (Fig. 6b-c). Moreover, most of the H3K4me3-only genes showed specific motif enrichment for the E-26 transformation specific (ETS) and nuclear transcription factor-Y (NF-Y) families, while the other genes with defect 2 were putative targets of homeodomain transcription factors, including HOX proteins (Fig. 6b).

**Figure 6:**
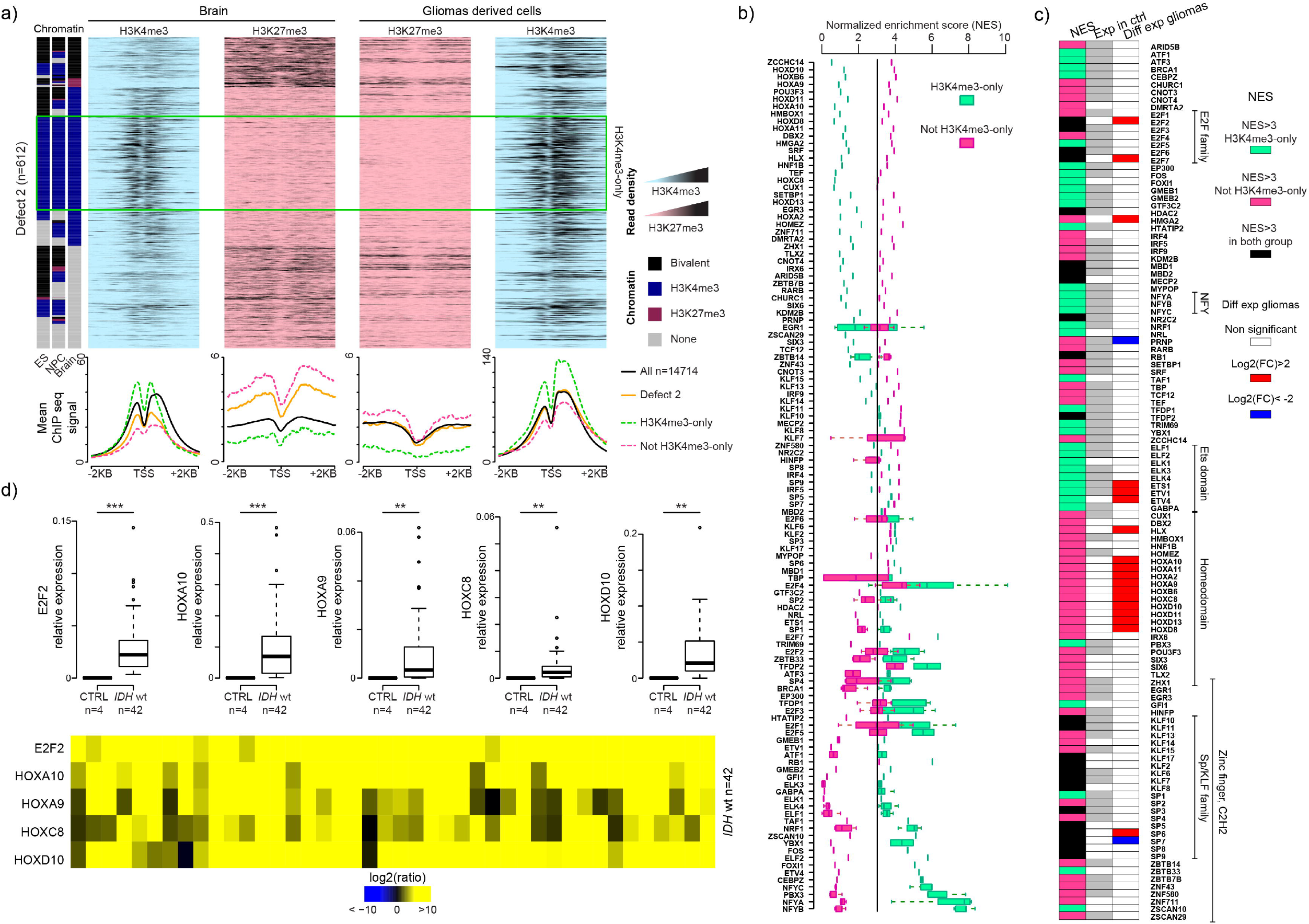
Transcription factor binding motifs in the promoters of genes overexpressed in glioma samples. **a)** Data-mining derived ChIP-seq read density data for H3K27me3 (purple) and H3K4me3 (blue) at genes with defect 2 in a ± 2Kbp window centered on their TSS, in healthy brain (left panels) and IDHwt-derived cell lines (right panels). The mean ChIP-seq signal values are shown on the lower panel for all class 2 defect genes (orange line) and for those that are (dotted green line) or not (dotted pink line) marked by H3K4me3-only in ES cells, NPC and brain. The black line, used as normalized reference, shows the value for all analyzed genes. **b)** Transcription factor motif enrichment in the CGI/promoter of class 2 defect genes, calculated using i-cisTarget and represented as a Normalized Enrichment Score (NES). Enrichment is shown for genes that are (green squares) or not (red squares) marked by H3K4me3-only in ES cells, NPC and brain. When a transcription factor possesses several binding motifs, data are presented as a box plot. **c)** Expression status, assessed by RNA-seq, of the transcription factors identified in b). The middle column shows their expression status in healthy control (n=5) (white, not expressed; gray, expressed: fpkm>1) and the right column their expression in IDHwt glioma samples (n=8). The left column shows the motif enrichment in all class 2 defect genes (black), and those with H3K4me3-only (green) and without (red) H3K4me3-only (red) in ES cells, NPC and brain. **d)** Expression *vs* controls of selected overexpressed transcription factor identified in (c) assessed by RT-qPCR in 42 IDHwt glioma samples. Details for each sample are provided in the lower panel (p-value by Mann-Whitney test).

Most of these transcription factors were expressed in healthy controls and remained expressed in glioma samples (Fig. 6c). However, a subset was specifically overexpressed in IDHwt glioma samples, including *HOX* genes, *E2F2* and *7, ETS1* and *4*, and *ETV4* (Fig. 6c: RNA-seq data). These findings were confirmed by RT-qPCR analysis of selected genes in 42 IDHwt samples (Fig. 6d). Gene encoding aberrantly overexpressed transcription factors mostly belonged to the defect 1 group (e.g., *HOXA2* and *HOXD8*) and defect 2 group (e.g., E2F2) (Additional file 2: Table S2). This observation suggests that the initial overexpression of few key transcription factors could lead to overexpression of most genes belonging to the defect 2 group.

### The repressive signature of silenced genes is related to their transcriptional status in healthy brain

To understand how genes with defect 3 or 4 were transcriptionally repressed, we compared their chromatin signature in healthy brain and in glioma samples using publicly available ChIP-seq data. This highlighted a marked enrichment for H3K27me3 in both groups in glioma samples compared with controls (Fig. 7a), but for the subset of genes with defect 4 that displayed the constitutive H3K4me3-only signature in ES cells, NPC and brain (framed in Fig. 7a) and did maintain this signature in glioma cells. This H3K27me3 gain in glioma cells led to a bivalent chromatin signature in the large subset of genes that were also marked by H3K4me3 (Fig. 7a). At CGIs/promoters of genes with defect 3 group, H3K4me3 tended to be reduced and H3K27me3 was enriched (Fig. 7a), suggesting that unlike normal cells (38), both H3K27me3 and DNA methylation can coexist at CGI/promoters in glioma cells.

**Figure 7:**
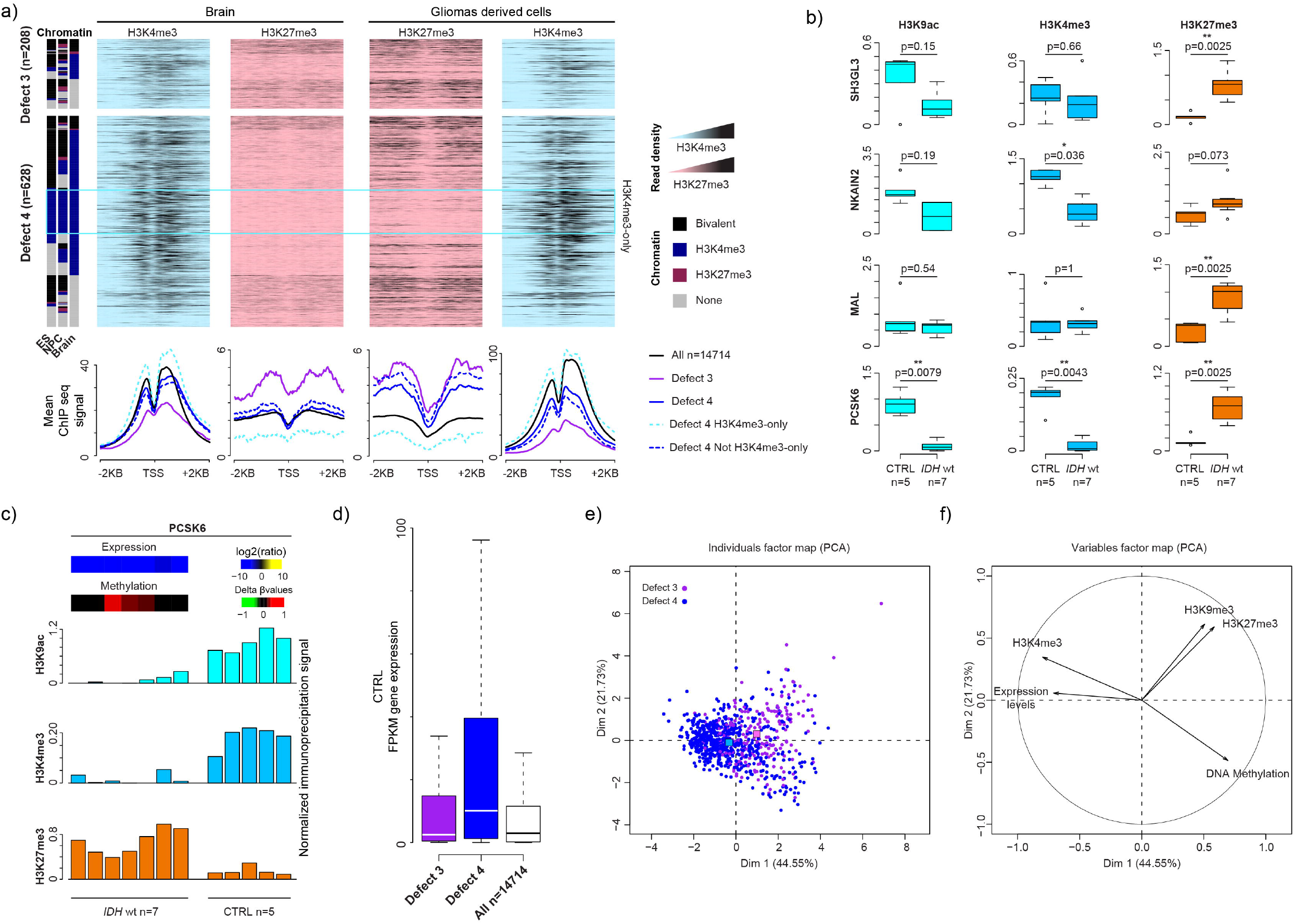
Gene repression is associated with H3K27me3 gain. **a)** Data-mining derived ChIP-seq read density data for H3K27me3 (purple) and H3K4me3 (blue) at genes with defect 3 or 4 in a ± 2 Kbp window centered on their TSS, in healthy brain (left panels) and IDHwt-derived cell lines (right panels). The mean ChIP-seq signal values are shown on the lower panel for class 3 (purple line) and class 4 (blue line) defect genes. Genes in class 4 were further subdivided in genes marked (dotted light blue line) and not marked (dotted dark blue line) by H3K4me3-only in ES cells, NPC and brain. The black line used as normalized reference shows the value for all analyzed genes. **b)** ChIP analysis of H3K9ac, H3K4me3 and H3K27me3 at selected genes in IDHwt (n=5) and control (n=7) samples. The precipitation level was normalized to that obtained at the *TBP* promoter (for H3K4me3 and H3K9ac) and at the *SP6* promoter (for H3K27me3) (p-values calculated with the Mann-Whitney test). **c)** Detail for each samples of the ChIP analysis at the *PCSK6* locus. Heat maps of the expression and methylation values are in the upper panel. **d)** Expression level of genes with defect 3 (purple column), 4 (blue column) and for all analyzed genes (white column) in healthy controls. **e-f)** Principal component analysis. Two-dimensional scatter plot of the values of each genes with defect 3 (purple dots) or 4 (blue dots) along the 1^st^ (Dim 1) and 2^nd^ (dim 2) principal components (e). For each class defect, the centroids are shown by colored squares. f) H3K4me3 and expression levels in healthy brain are the variables that most contributed to and were significantly correlated with the first principal component.

ChIP analysis of selected genes from the defect 4 group (*PCSK6, MAL, SH3GL3* and *NKAIN2*) confirmed the marked H3K27me3 gain associated with a decrease or maintenance of H3K4me3 and H3K9ac, according the studied locus, in the seven glioma samples tested (Fig. 7b-c, and Additional file 1: Figure S5a). Also, bisulfite analysis of the H3K27me3-immunopreciptated fraction confirmed that both DNA methylation and H3K27me3 co-existed at the CGI/promoter of genes with defect 3 in glioma samples, as exemplified by the methylable *PCSK6* gene (Additional file 1: Figure S5b).

Several studies have highlighted that the propensity of genes to be hypermethylated in cancer cells is related to their transcriptional status in the normal tissue (7–11). Accordingly, we observed that the transcriptional status in brain discriminated between genes in the defect 3 and 4 groups. Specifically, DNA methylation-independent silencing (defect 4) mainly affected genes that were highly expressed in brain. Conversely, poorly expressed genes tended to be hypermethylated (Fig. 7d). To more systematically assess the bases of variation between these two groups, we performed principal component analysis for all genes in the defect 3 and 4 groups. The first principal component accounted for about 44.5% of the variance and allowed separating the two defect groups (centroid values along the axis: +0.984 and −0.326, respectively) (Fig. 7e). The observation that the first principal component was significantly correlated with the permissive H3K4me3 mark (R=0.79; P < 3.4×10^−126^) and the expression (R=0.71; P < 1 x10^−129^) levels in healthy brain (Fig. 7f) supported the hypothesis that the expression status in healthy cells contribute to the choice of silencing pathway used in cancer cells. Additional analyses using normalized RNA-seq data for 21 different tissues showed that genes from the defect 4 groups were strongly expressed specifically in adult brain tissues (Additional file 1: Figure S5c and d).

Thus, besides DNA methylation, gain of H3K27me3 and chromatin bivalency emerges as major causes of aberrant gene repression in aggressive glioma cells. The choice between these alternative silencing pathways is related to the expression level of the affected genes in healthy tissues/cells.

## Discussion

Here we used glioma, one the most widespread brain tumor types, to evaluate the relative contribution of DNA methylation-dependent and -independent mechanisms to transcriptional alteration at CGI/promoter-associated genes in cancer cells. Our study highlighted that H3K27me3 level changes are the predominant molecular defect at both aberrantly repressed and expressed genes. Moreover, our findings support that altered control of H3K27me3 dynamics, more specifically when present in a bivalent context, is the main cause of transcriptional alteration in glioma cells.

A handful of studies have reported H3K27me3-based transcriptional repression in cancer cells (7, 18–20, 39). In colorectal tumors, ectopic gene expression has been associated with aberrant loss of H3K27me3 from CGI/promoters with bivalent chromatin signature (20). Moreover, gene expression changes in primary human clear cell renal cell carcinomas can be attributed to chromatin accessibility alterations, independently of DNA methylation (40).

Our study further extends these observations and provides for the first time a comprehensive description of these alterations in glioma samples and their relative contribution to transcriptional alteration in such tumors. Specifically, our integrative analyses identified and quantified four main types of transcriptional defects in glioma (Fig 8) that recapitulate the DNA methylation- and H3K27me3-based molecular signatures at aberrantly repressed and expressed CGI/promoter-associated genes. We detected these defects in IDHwt and also IDHmut glioma samples (Fig 8b, Additional file 1: Figures S6 & S7), indicating that they occur regardless of the G-CIMP status and clinical outcome, and suggesting that our observations might apply to cancer cells in general.

**Figure 8:**
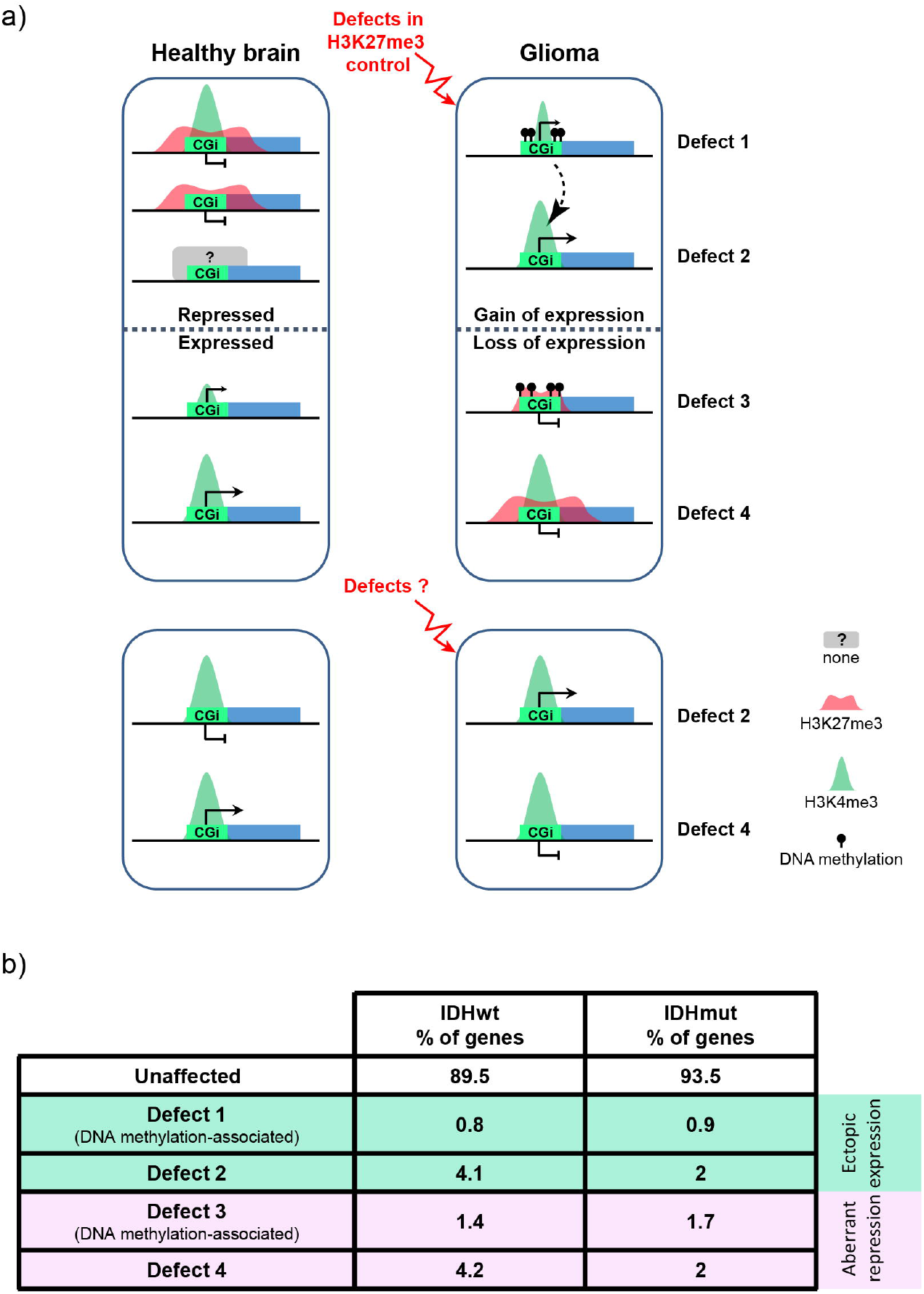
Working model. **a)** In glioma, alterations in the control of the H3K27me3 signature could be one of the main contributors to the four types of transcriptional defects observed at CGI/promoter-controlled genes (upper panels). In this model, genome-wide hypomethylation induces H3K27me3 redistribution that could lead to ectopic expression of genes that are normally repressed by polycomb proteins, including some genes encoding transcription factors. These overexpressed transcription factors could then promote the aberrant expression of their target genes (dotted arrow). Similarly, alterations in the interplay between the polycomb complex and the transcriptional machinery could affect H3K27me3 fate during ES and/or neural stem cell differentiation. Specifically, this alteration could lead to the aberrant maintenance of bivalency and silencing at a subset of genes that are normally specifically expressed in brain. At genes that are normally poorly expressed in healthy brain, this process is associated with gain of DNA methylation in glioma. Beside defects in the H3K27me3 signature, we also identified a subset of genes that are apparently constitutively associated with H3K4me3-only, regardless of their expression status in brain and glioma (lower panel). The mechanisms underlying their transcriptional deregulation remain to be determined. **b)** Percentage of unaffected and affected CGI/promoter-controlled genes for each of the four defects described in the article, in IDHwt and IDHmut glioma samples.

Our study revisited the relationship between aberrant transcription and DNA methylation in cancer cells. We first confirmed, in agreement with the general unmethylated status of CGI/promoters in healthy cells, that gene expression deregulation is very rarely associated with CGI/promoter DNA hypomethylation in glioma samples. Unexpectedly, we revealed that DNA hypermethylation is not the main cause of transcriptional repression at CGI/promoter genes. In addition, it can be associated with gain of expression. For instance, about 16% of ectopically expressed genes were associated with a hypermethylated CGI/promoter in IDHwt glioma samples. Specifically, in many genes, ectopic expression was associated with CGI/promoters that gained methylation at their borders, while the H3K4me3-marked TSS was methylation-free. At other genes, extensive methylation of their main CGI/promoter was associated with the use of an alternative promoter. Interestingly, these two distinct signatures have also been described in prostate cancer cell lines (21), suggesting that an association between CGI/promoter DNA hypermethylation and gain of gene expression is common in cancer. Besides the concomitant gain of expression and DNA methylation, this group of genes was characterized also by a dramatic reduction of H3K27me3 level in glioma cells (compared with controls), suggesting that the interplay between these two repressive marks is a driving force in their transcriptional deregulation. Specifically, studies in the mouse highlighted that widespread DNA methylation depletion triggers redistribution of H3K27me3 (38, 41) that in turn leads to a dramatic loss of H3K27me3 and ectopic expression at a subset of polycomb target genes, including *Hox* clusters (41). Reciprocally, analyses in mouse ES cells showed that vicinity of CGI/promoters of polycomb target genes is protected from aberrant gain of DNA methylation by polycomb proteins (42). We thus propose that defect 1 affects a subset of polycomb-target genes that are particularly sensitive to the H3K27me3 redistribution induced by the genome-wide hypomethylation of cancer cells. It is noteworthy that in the aggressive IDHwt glioma, the defect 1 group is enriched in homeodomain genes, and more specifically in *HOX* genes. It has been shown that deregulation of *HOX* genes contributes to the tumorigenic potential of glioblastoma stem cells, by activating a network of downstream genes (43). Accordingly, we observed that many ectopically expressed genes in the defect 2 groups are putative targets of HOX transcription factors. Altogether, our observations support a domino effect model to account for the gain of expression of CGI/promoter genes in aggressive glioma. In this model, genome-wide hypomethylation leads to ectopic expression (and methylation gain) of genes in the defect 1 group (especially *HOX* genes) than then promote gain of expression of target genes with defect 2 (Fig. 8a). Additional studies are required to test this model and specifically whether genes with defect 2 are *bona fide* targets of HOX transcription factors.

Another key finding of our study is that a bivalent chromatin signature in stem cells may not only predispose genes to hypermethylation, as widely documented (11, 32, 33), but more globally to transcriptional alteration in cancer cells. We observed that genes with CGI/promoters marked by a bivalent chromatin signature in ES cells and NPC are more prone to be deregulated in glioma samples, regardless of the type of transcriptional defect. This was particularly true for genes with DNA methylation-associated defects, irrespectively of their association to gain or loss of expression (defect 1 and 3), and to a lesser extent, for genes with DNA methylation-independent defects (groups 2 and 4). This observation suggests that defects in the control of the bivalent chromatin signature, and more specifically of H3K27me3 dynamics, upon differentiation, are one of the main causes of transcriptional deregulation of CGI/genes in cancer cells. Accordingly, we observed that aberrant repression in glioma samples affected mainly genes with brain-specific expression and thus more sensitive to bivalency alterations upon neural stem cell differentiation. Besides the functional aberrations of writers and erasers of the H3K27me3 and H3K4me3 marks documented in many tumors (16,17), these control defects could also result from transcriptional changes in key tissue-specific transcription factors or co-factors. Studies in mouse ES cells and tissues highlighted that their transcriptionally inactive status is sufficient to promote the recruitment of the polycomb responsive complex PRC2 and H3K27me3 deposition at CGI/promoters (44–48). This suggests that the fate of bivalent chromatin domains upon development/differentiation relies on the interplay between PRC2 and the availability of the *ad hoc* transcriptional machinery. Our observation that the gene transcription strength in healthy brain influences the choice of silencing mechanism at repressed genes in glioma precisely argues for an alteration in this interplay. Specifically, we propose that following the alteration of a subset of brain-specific co-factors, the resulting weakened transcriptional machinery cannot efficiently counteract PRC2 recruitment upon differentiation, leading to aberrant maintenance of chromatin bivalency and to silencing of a subset of genes that are normally specifically expressed in brain. Moreover, gain of function of factors that promote, directly or indirectly, the recruitment of PRC2 at CGI could facilitate this process. This includes for instance the histone demethylase KDM2B (49–51) that is critical in various cancers, including glioma (52,53). At normally poorly expressed genes, these events associated with the initial weak level of H3K4me3, a mark that prevents recruitment of DNA methyltransferases, would facilitate the subsequent gain of DNA methylation (Fig. 8a).

In addition to genes in which their chromatin signature was altered in glioma, we also identified a subset of genes with an apparent constitutive H3K4me3-only signature, detected in healthy (ES cells to brain) and glioma samples, and that showed either gain of expression (defect 2) or aberrant repression (defect 4) in glioma samples (Fig. 8a). Additional studies are required to establish the molecular bases of these observations. Specifically, it would be interesting to determine whether the regulation of these genes in normal and pathological contexts relies exclusively on the availability of *ad hoc* transcription factor(s), or whether it is also associated with not yet identified repressive histone marks. Moreover, genes in this group that are ectopically expressed in glioma samples were also expressed in ES cells and NPC (Fig. 4). Similarly to genes with bivalent chromatin signature in ES cells whose aberrant repression in glioma also recapitulated their repression in stem cells (Fig. 4), this group of genes could contribute to glioma aggressiveness by maintaining tumor cells in a stem-cell like state.

Our study also provides a framework to explain the counterintuitive observation that patients with CIMP-positive IDHmut have a better clinical outcome than patients with CIMP-negative IDHwt. Our data indicate that CIMP is observed mainly at genes that are already repressed in healthy brain. Consequently, the number of deregulated genes with CGI/promoters associated with DNA methylation defects is similar between glioma subtypes. Conversely, the higher frequency (about two times) of DNA methylation-independent transcriptional alterations in IDHwt than in IDHmut samples could contribute to the prognosis difference between glioma subtypes. The CIMP-positive status, by promoting stable gene repression, could also act as a protective mechanism against cancer progression, by limiting the tumor epigenetic plasticity and its ability to adapt to environmental changes, such as metastasis formation or treatment (54). Specifically, among the hundreds of genes that gain expression in IDHwt samples and that are maintained repressed through gain of methylation in IDHmut samples (Additional file 4: Table S4), a dozen are oncogenes with some documented for their role in glioma biology, such as Spalt-like transcription factor 4 (*SALL4*) (55) and the long non-coding RNA MIR155 host gene (*MIR155HG*) (56).

## Conclusion

Our study on the extent and consequences of epigenetic alterations in glioma indicates that transcriptional deregulations rely mainly on chromatin-based DNA methylation-independent mechanisms. It also shows that the gene expression level in healthy tissue influences the type of silencing pathway used for repression in cancer cells, whereby highly expressed genes are more likely to be repressed by H3K27me3 rather than DNA methylation. Besides providing an original framework to understand the epigenetic basis of carcinogenesis, these observations are also important for the design of drugs to target epigenetic defects in cancer.

## Materials and Methods

### Tumor and control samples

Glioma samples (*n*=70) were from tumors resected during the standard diagnostic and treatment procedure. Immediately after surgery, samples were snap-frozen and stored in liquid nitrogen. Random sections of each tumor were analyzed under a light microscope after hematoxylin–eosin staining to determine the extent of necrosis and the percentage of tumor cells. All glioma samples included more than 50% of tumor cells. Gliomas were classified according their IDH1 mutation status: IDHwt (n=55) and IDHmut (n=15). *IDH1* genotyping was performed by EpigenDx (Worcester, MA) (EpigenDx pyrosequencing assays ADS1703 and ADS1704) (57).

Eight control brain samples (healthy controls, mean age of 27.3 years, standard deviation ± 2 years) were removed by autopsy 4-16h after accidental death (Brain and Tissue Bank of Maryland). These samples, identified by the Brain and Tissue Bank of Maryland as corpus callosum (n = 5) and frontal cortex (n = 3), corresponded to white matter enriched in astrocytes and oligodendrocytes from which gliomas originate.

Tumor and control samples were homogenized into powder by cryogenic grinding, equally distributed in at least three vials (57) before use for matched genomic DNA, RNA and chromatin extraction. All samples were stored at −80°C until use. Overall survival (OS) was calculated as the number of days between the surgery date and the patient’s death. Tumor resection was classified as gross total resection, when no enhanced contrast was detected 48h post-surgery, or as partial resection, when enhanced contrast was still detected 48h post-surgery.

The demographic and clinical features are presented in Additional file 1: Table S1. The related statistical analyses were performed using the Stata software, version 13 (StataCorp, College Station, TX, US) and the R software, version 3.3.1. To test the prognostic value of the patients’ characteristics in univariate analyses, OS curves were compared between groups using the log-rank test. Results are expressed as hazard ratios (HRs) and 95% confidence intervals (CIs).

### Selection of the genes to be analyzed

The positions of genes and CpG islands (CGI: defined using standard criteria: *GC content* ≥*50%; length* >*200bp; ratio Obs/Exp CpG* >*0.6*) were downloaded from Gencode annotation release V19, and CpG-island tracks of UCSC hg19 assembly, respectively. For each gene, the promoter region was defined as TSS ± 1kbp. To assess the relationship between CGI/promoter methylation status and gene expression, we first identified the genes associated with only one promoter with CGI features (n=15350). As our cohort included both men and women, we then excluded genes located on the X or Y chromosomes. We finally retained 14714 genes in which the CGI/promoter is covered by at least two probes in the HM450K Illumina array.

### DNA methylation analyses

#### DNA extraction

DNA was isolated from frozen tissue samples using the QIAamp DNA Mini Kit (Qiagen, Hamburg, GmbH) according to the manufacturer’s recommendations.

#### Gene-specific bisulfite sequencing

Bisulfite conversion, PCR amplification, cloning, and sequencing were performed as previously described (58). Details on the primers used are in Additional file 5: Table S5.

#### HM450K analysis

The HM450K array data for controls and gliomas sample (8 normal brain, 55 IDHwt and 15 IDHmut gliomas) were analyzed as previously described (59). Specifically, DNA bisulfite conversion and array hybridization were performed by Integragen SA (Evry, France), using the Illumina Infinium HD methylation protocol. β-values were computed using the GenomeStudio control interplate normalization and background subtraction (version 2011.1, manifest files: HumanMethylation450_1 5017482_v.1.2.bpm). For each sample, β-values with a detection P-value >0.01 were excluded. All probes with a detection P-value >0.01or lacking signal in more than 5% of our samples were excluded. Finally, 26 507 probes containing common SNPs (dbSNP 147) in their last 5bp or in the CpG sites were discarded. As our patient cohort included both men and women, probes on the X and Y chromosomes were also excluded from the analysis. After the implementation of these quality filters, a total of 443 691 CpG methylation values were considered suitable for the downstream analysis. Methylation level at the 14714 genes (11795 CGIs) was given by the mean β-value of all probes located in their CGI. Differential methylation analyses were performed using the limma R package, as previously described (60). These analyses concerned the entire groups (8 controls vs 55 IDHwt, and 8 controls *vs* 15 IDHmut) or only the samples with matched RNA-seq data (3 controls *vs* 8 IDHwt and 3 controls vs 5 IDHmut). The HM450K probes were considered differentially methylated when the FDR was <0.05 and when the β-value difference between groups was >0.1. To consider only robust methylation variations, CGI/promoters were classified as hyper-methylated or hypo-methylated only if they included at least two probes differentially methylated (gain or a loss of methylation) in their CGI.

### Expression analysis

#### RNA extraction

RNA was isolated from frozen tissue samples using the RNeasy Mini Kit (Qiagen, Hamburg, GmbH), according to the manufacturer’s recommendations.

#### RT-qPCR expression analyses

Rt-qPCR assays were performed using a microfluidic-based approach, as previously described (48). First-strand cDNA was pre-amplified for 14 cycles with the pool of primers used for RT-qPCR and the Taq-Man PreAmplification Master Mix (Life Technologies, 4488593). Primer sequences are in Additional file 5: Table S5. RT-qPCR assays were then performed and validated using Fluidigm 96.96 Dynamic Arrays and the Biomark HD system (Fluidigm Corp.), according to the manufacturer’s instructions. The relative expression level was quantified with the 2-dCt (Livak et al, 2001). The housekeeping genes PPIA, *TBP* and *RPL13A* were used to normalize transcript expression. Each analysis was performed in duplicate.

#### RNA-sequencing

RNA-seq was performed using total RNA after ribosomal RNA depletion (Ribo-Zero rRNA Removal Kit, Illumina) from 3 brain, 8 IDHwt and 5 IDHmut glioma samples. RNA processing, rRNA depletion, library preparation and sequencing on an Illumina Hiseq 2500 apparatus were performed by Integragen SA (Evry, France), according to the manufacturer’s protocol (mean of 90 million of paired reads per sample). Stranded RNA-seq reads were mapped to the human genome (hg19) using tophat2 (version 2.1.0) and a transcript annotation file from Gencode (Release 19). The average alignment rate was about 94.5% with a concordant pair alignment rate of 92%. Only properly paired reads were considered for downstream analysis. The reads count per gene was obtained with the HTseq-count script (option: -m intersection-nonempty -s reverse), and the FPKM gene expression level was determined with Cuffquant and Cuffnorm from the Cufflinks suite (version 2.2.1), based on Genecode V19 transcript annotation. Strand-specific RNA-seq signals were derived from the RNA-seq alignments using samtools, genomeCoverageBed and bedGraphToBigWig tools, and visualized using the UCSC Genome browser. Differential expression analyses between controls and glioma samples were based on reads counts using the DESeq2 and edgeR R packages. Genes were considered as differentially expressed between groups when |log_2_-fold change| >2 with an adjusted P-value <0.05 in both statistical approaches.

#### Gene expression data mining

Gene expression levels in several human tissues were obtained from the Roadmap Epigenomics project (https://egg2.wustl.edu/roadmap/web_portal/processed_data.html#RNAseq_uni_proc). The transcription levels in different brain regions were retrieved from the GEO database (accession number GSE33587).

### Chromatin analyses

#### Chromatin immuno-precipitation from glioma samples

Anti-H3K9ac (Millipore 06-942), -H3K4me3 (Diagenode 03-050) and -H3K27me3 (Millipore 07-449) antibodies were used to assess the respective marks at selected genes by ChIP of native chromatin isolated from glioma samples and controls, as described in (48). Input and antibody-bound fractions were quantified by real-time PCR amplification with the SYBR Green mixture (Roche) using a LightCycler 480II (Roche) instrument. Background precipitation levels were determined by performing mock precipitations with a non-specific IgG antiserum (Sigma C-2288) and were only a fraction of the precipitation levels obtained with the specific antibodies. The bound/input ratios were calculated and normalized to the precipitation level at the *TBP* promoter for the anti-H3K9ac and - H3K4me3 ChIPs and at the *SP6* promoter for the anti-H3K27me3 ChIP. The primers used are described in Additional file 5: Table S5.

#### ChIP-seq data mining associated with chromatin analyses

ChIP-seq data from NPC and brain samples were from the NIH Roadmap Epigenomics project (http://www.roadmapepigenomics.org/). In detail:

- For H9-derived Neural Progenitor Cells (NPC): Input (GSM772805); H3K4me3 (GSM772736), and H3K27me3 (GSM772801)
- In brain: Input (GSM772991), H3K4me3 (GSM772996), H3K27me3 (GSM772993) and H3K9me3 (GSM670005)

ChIP-seq data for the H3K4me3 and H3K27me3 profiles in glioma-derived cells were obtained from the GEO database (accession numbers: GSM1121888 and GSM1121885, respectively). To describe the histone modification enrichment in each defect group, the ChIP-seq read densities around TSS (± 2 Kb) was represented by a heatmap where each line represents one single promoter. The mean signal around TSS (± 2 Kb) for each defect group was compared with the mean signal for all genes included in this study (n=14714) to correct for the bias due to the use of different datasets.

#### Gene classification according to their chromatin signature

The gene classification according to their chromatin signature (bivalent, H3K4me3-only, H3K27me3-only and none) in human ES cells is from (11). For NPC and brain samples, this classification was performed as previously described in (11). Briefly, data for input, H3K4me3 and H3K27me3 ChIPs were aligned to the hg19 genome assembly. Peaks were then called with Macs 1.4.2 using the input as control for peak detection. Chromatin signatures were classified using an in-house R scripts. Specifically, a bivalent region was defined by the overlapping of H3K4me3 and H3K27me3 peaks for at least 1Kbp. H3K4me3-only and H3K27me3-only regions were identified as 1Kbp peaks for H3K4me3 or H3K27me3 that did not overlap. The rest of the genome was considered as having a “none” chromatin signature. To attribute a chromatin signature to CGI/promoter regions, only the signature spreading across the CGI region was retained.

### Copy-number variation (CNV) analyses

CNV analyses were performed using the Genome-Wide Human CytoScan HD Array (Affymetrix, CA, USA) and the samples analyzed by RNA-seq (3 controls, 8 IDHwt and 5 IDHmut glioma samples). DNA and array data were processed by the Genomic Platform/Curie Institute (France) according to the manufacturer’s protocol. Arrays were scanned using an Affymetrix GeneChip Scanner 3000 7G. Scanned data files were analyzed with Affymetrix Chromosome Analysis Suite v3.1 (ChAS) (Affymetrix Inc., USA) using the CytoScanHD_Array.na33.annot.db annotation file on the hg19 genome. To find CNVs, single samples were analyzed using the reference model file “CytoScanHD_Array.na33.r2.REF_MODEL”. To prevent the detection of false-positive CNVs, only alterations that involved at least 50 consecutive probes and spanning more than 100 kbp were considered in the analysis. To evaluate genetic alterations at gene level, the mean log2 ratio of the CNV fragment overlapping with a CGI/promoter region was assigned to each gene. For genes with a CGI/promoter region not covered by a CNV fragment, the mean log2 ratio was considered as null. Positive and negative mean log2 ratios were used to categorize duplicated and deleted regions, respectively.

To identify genes the expression alteration of which correlates (p <0.05) with CNV changes, the mean log2 ratio of the CNV values and the normalized pseudo RNA-seq counts for each transcriptionally affected gene in the same sample were compared using the for the Pearson correlation.

### Principal component analysis (PCA)

The PCA was done with the FactoMineR package using molecular features (histones modifications, DNA methylation and expression) associated with genes from the defect 3 and 4 groups in brain. For each gene, the histone modification levels at the CGI/promoter region (TSS ±1kb) were based on the average ChIP-seq signal for H3K4me3, H3K27me3 and H3K9me3 obtained in brain samples. The DNA methylation levels were determined by the log2 value of the mean β-value, obtained from 8 brain control samples, for all the probes located in the CGI of that gene. For the transcriptional level, the average FPKM value from 3 brain control samples was log2 transformed. As the different variables used were defined by different measure units, they were standardized all before the PCA analysis (i.e., centered and scaled).

### Functional annotations

InterPro protein functional classification analysis was performed using the functional annotation tools in DAVID 6.8 (https://david.ncifcrf.gov/). Gene ontology analyses were done with the GeneRanker tools of the Genomatix suite (http://www.genomatix.de/). To identify putative regulatory features linked to the transcriptional defect groups, the CGI positions in genes were analyzed with *i-cis* Target (https://gbiomed.kuleuven.be/apps/lcb/i-cisTarget/). The comparative analysis tools were then used to identify specific binding sites. As transcription factors could have multiple position weight matrices (PWM), all Normalized Enrichment Scores (NES) given by *i-cis* Tagert for a factor were displayed in a boxplot. The tumor-suppressor gene and oncogene lists were obtained from www.ongene.bioinfo-minzhao.org, www.cta.lncc.br and www.uniprot.org with the keywords tumor suppressor (KW-0043) and proto-oncogene (KW-0656).

## Abbreviations

CGI: CpG island
ChIP: chromatin immunoprecipitation
CI: Confidence interval
CIMP: CpG island methylator phenotype
CNV: Copy number variation
EZH2: Enhancer of Zeste homolog 2
FPKM: Fragments per kilobase of transcript per million mapped reads
G-CIMP: Glioma-CIMP
HR: Hazard ration
IDH: Isocitrate dehydrogenase
IDHmut: IDH mutated
IDHwt: IDH wild-type
INA: Alpha-internexin
MLL: Mixed-lineage leukemia
NPC: Neural Progenitor Cells
PCA: principal component analysis
RNA-Seq: RNA sequencing
TSS: transcription start site
UTX: Ubiquitously transcribed tetratricopeptide repeat, X chromosome.

## Declarations

### Ethics approval and consent to participate

Glioma samples (*n*=70) resected between 2007 and 2014 were obtained from Clermont-Ferrand University Hospital Center, France (Anonymized samples from the “Tumorotheque Auvergne Gliomes’, ethical approval DC-2012-1584). The ethics committees and the respective competent authorities approved this study. The study protocols conform to the World Medical Association Declaration of Helsinki.

### Consent for publication

“Not applicable”

### Availability of data and materials

Data generated in this study have been submitted to the NCBI Gene Expression Omnibus (GEO; http://www.ncbi.nlm.nih.gov/geo/) under the accession numbers:

- GSE123678 for the the HM450K DNA methylation data
- GSE123682 for the Cytoscan HD data

GSE123892 for the oriented RNA-seq data ChIP-seq data from NPC and brain were obtained from the NIH Roadmap Epigenomics project (http://www.roadmapepigenomics.org/). In detail:

- H9-derived Neural Progenitor Cells (NPC): Input (GSM772805); H3K4me3 (GSM772736). H3K27me3 (GSM772801)
- Brain: Input (GSM772991); H3K4me3 (GSM772996). H3K27me3 (GSM772993) H3K9me3 (GSM670005) ChIP-seq data for the H3K4me3 and H3K27me3 profiles in gliomas derived cells were obtained from the GEO database (accession numbers: GSM1121888 and GSM1121885, respectively).

### Authors’ contributions

FC and PA initiated and supervised the study. FC, AF, CVB, EC, LKT, PV and PA designed the study. TK collected glioma samples from the operating room. JLK, JB, EC and PV characterized the glioma samples and updated the clinical follow-up of the patients. JB and EC collected the patients’ data. BP conducted the patients’ data statistical analyses. FC, ELB, AF, MMB and CVB performed the experiments. FC performed the bioinformatics analyses. FC, ELB, AF, MMB, CVB, EC and PA analyzed the data. FC produced the figures with CVB and PA input. PA wrote the paper with FC input. All authors read and approved the final manuscript.

## Acknowledgments

We thank the CHU Clermont-Ferrand’s Neurosurgery department (Prof. JJ. Lemaire) for its help in completing this study, the Platform “Gentyane” (http://gentyane.clermont.inra.fr/) for its technical help with the Fluidigm 96.96 Dynamic Arrays, and Zachary-LoL-Arnaud for its precious input. We also thank Dr David Monk and all members of PA’s team for critical reading of the manuscript.

## Additional Files

**Additional file 1: Supplementary Table S1 & Supplementary Figures S1 to S7 (PDF)**

- **Table S1:** Demographic and molecular features of the 70 patients with glioma
- **Figure S1: DNA methylation alteration in IDHwt and IDHmut glioma samples**
- **Figure S2: Transcriptional alterations in IDHwt and IDHmut glioma samples**.
- **Figure S3: Gene ontology**
- **Figure S4: Expression of genes with a methylated CGI/promoter**
- **Figure S5: Gene repression is associated with H3K27me3 gain and affect mainly genes with brain-specific expression profile**
- **Figure S6: Four classes of expression defects in IDHmut glioma samples**
- **Figure S7: Genes with bivalent chromatin signature in ES cells are more prone to be deregulated in IDHmut glioma.**

**Additional file 2: Supplementary Table S2 (XLSX)**

List of CGI/promoter-controlled genes with defect 1 to 4, identified in IDHwt and IDHmut glioma samples.

**Additional file 3: Supplementary Table S3 (XLSX)**

List of CGI/promoter-controlled genes in which expression alteration correlated with copy number changes in IDHwt and IDHmut glioma samples.

**Additional file 4: Supplementary Table S4 (XLSX)**

List of genes that are upregulated in IDHwt glioma samples and that are maintained repressed with a gain of methylation in IDHmut glioma samples.

**Additional file 5: Supplementary Table S5 (XLSX)**

Primer list

**Figure S1: DNA methylation alteration in IDHwt and IDHmut glioma samples**

**a)** Kaplan-Meier survival curves for patients with IDHwt (n=55) and IDHmut (n=15) glioma confirmed that the IDH status is a significant prognostic marker: 2-year survival rate of 27% in patients with IDHwt (median survival of 1.2 years/14.4 months) and 83% in patients with IDHmut glioma (median survival not reached). **b)** Visualization of the DNA methylation level (β-values) at the 86157 probes (row) located in the 11795 CGI/promoters analyzed in this study in IDHmut, controls and IDHwt glioma samples (column) identified a G-CIMP profile in IDHmut glioma samples. **c)** Differential DNA methylation level vs controls (delta of the means of β-values) of the 11795 CGIs (row) analyzed in this study in IDHmut and IDHwt glioma samples (columns). **d)** Classification of the genes associated with hyper-, hypo-methylated or unaffected CGI/promoters, respectively, in IDHwt (upper panel) and IDHmut (lower panel) glioma samples.

**Figure S2: Transcriptional alterations in IDHwt and IDHmut glioma samples.**

**a)** Volcano plot analysis with DESeq2 of the differential expression of all annotated genes (n= 44267) in IDHwt (left) and IDHmut (right) glioma samples compared with healthy brain controls. Genes that were significantly up- or down-regulated are symbolized by a red dot for CGI/promoter-associated genes, and by a black dot for the others. The numerical values are given in the table. **b)** Distribution per chromosome of transcriptionally up- (upper panels) and down-regulated (lower panels) CGI/promoter-associated genes in IDHwt and IDHmut glioma samples. The relative proportion of gene in which transcription alteration correlated with CNV is indicated. **c)** Details of the correlation analyses between CNV and expression for the *TTC18, EZH2*and *RBM42* genes in IDHwt (yellow dots in the left panels) and IDHmut (blue dots in the right panels) glioma samples. Black dots indicate the expression value in healthy brain. *TTC18* downregulation and *EZH2* overexpression correlated with CNV in IDHwt glioma samples. *RBM42* downregulation correlated with CNV in IDHmut glioma samples.

**Figure S3: Gene ontology**

**a)** Gene ontology terms enriched in genes with defect 1 to 4 identified in IDHwt glioma samples. For each category, the four highest terms are shown. **b)** Distribution of genes with defect 1 to 4 according to their chromatin signature in human NPC (none: gray; bivalent: black; H3K4me3-only: blue; H3K27me3-only: red). As reference, the distribution of the 14714 genes analyzed in this study according to their chromatin signatures in human ES cells is shown in the left panel.

**Figure S4: Expression of genes with a methylated CGI/promoter**

Genome browser view showing H3K4me3 enrichment, differential DNA methylation, and oriented RNA-seq signal at **a)** *CTHRC1* (representative example of genes in which transcription initiates from an H3K4me3-marked TSS embedded in a methylated CGI/promoter, **b)** *NR2F2* (example of genes in which transcription initiates from an alternative TSS, and **c)** *HEYL* and *C15orf44*, where transcription initiates from a methylated CGI/promoter.

**Figure S5: Gene repression is associated with H3K27me3 gain and affect mainly genes with brain-specific expression profile**

**a)** ChIP analysis of H3K9ac, H3K4me3 and H3K27me3 at selected genes in IDHwt glioma (n=7) and control (n=5) samples. The precipitation level was normalized to that obtained at the *TBP* promoter (for H3K4me3 and H3K9ac) and at the *SP6* promoter (for H3K27me3). **b)** Bisulfite-based sequencing data for the *PCSK6* CGI/promoter using input and H3K27me3-immunoprocipated ChIP fractions showed that DNA methylation and H3K27me3 can coexist in glioma cells. Each horizontal row of circles represents the CpG dinucleotides on an individual chromosome. Solid circles, methylated CpG dinucleotides; open circles, unmethylated CpG dinucleotides. The relative position of the bisulfite amplicon is showed on the *PCSK6* locus browser view (right panel). **c)** Median expression of genes with defect 3 (purple column), 4 (blue column) or all CGI/promoter genes (white column) in 21 tissues (publicly available normalized RNA-seq data). Genes with defect 4 are strongly expressed specifically in adult hippocampus **d)** The expression level in hippocampus is representative of the expression level in other brain parts, as shown for the frontal pole and caudal nucleus.

**Figure S6: Four classes of expression defects in IDHmut glioma samples**

**a)** Integrative analysis of gene expression and methylation assessed in 5 IDHmut glioma samples identified four main defect classes. **a)** Differential DNA methylation analysis of all the IDHmut glioma samples (n=15) classified according the defect classes defined in a). The methylated and methylable status of the genes is indicated in the left column. **c)** Integrative analysis of the differential expression and methylation (vs control) for selected genes from each class defect in 10 IDHmut glioma samples.

**Figure S7: Genes with bivalent chromatin signature in ES cells are more prone to be deregulated in IDHmut glioma.**

**a)** Gene ontology terms enriched in genes with defect 1 to 4 identified in IDHmut glioma samples. For each category, the four highest terms are shown. **b)** Distribution of genes with defects 1 to 4 according to their chromatin signatures in human ES cells (none: gray; bivalent: black; H3K4me3-only: blue; H3K27me3-only: red). As a reference, the distribution according to their chromatin signatures in human ES cells of all the 14714 genes analyzed in this study is shown in the left panel. **c)** Expression level and associated chromatin signatures of genes with defects 1 to 4 in human ES cells, neural progenitors cells (NPC) and healthy brains.

