## supplementary data for "Most of transcriptional alterations in glioma result from DNA-methylation independent mechanisms"

Supplementary Table 1: Demographic and molecular features of the 70 patients with glioma

| Characteristics | Mutated IDH1 patients (n=15) | | WT IDH1 patients (n=55) | |
|---|---|---|---|---|
| | Nb of patients | % | Nb of patients | % |
| Sex | | | | |
| Female | 5/15 | 33% | 23/55 | 42% |
| Male | 10/15 | 67% | 32/55 | 58% |
| Age at diagnosis (years) | | | | |
| Mean (SD) | 40 (10.4) | | 60 (10.2) | |
| Range | [26-63] | | [29-80] | |
| 7p+/10q- (CNV) | 0/13 | 0% | 23/34 | 68% |
| NA status 7p10q | 2/15 | 13% | 21/55 | 38% |
| Tot 1p19q codeleted (FISH+INA+CNV) | 9/14 | 64% | 2/55 | 4% |
| NA status 1p19q | 1/15 | 7% | 0/55 | 0% |
| *MGMT* promoter methylation (>8%) | 14/15 | 93% | 25/55 | 45% |
| Resection | | | | |
| Total | 6/15 | 40% | 23/55 | 42% |
| Partial | 7/15 | 47% | 31/55 | 56% |
| Large biopsy | 2/15 | 13% | 1/55 | 2% |
| Post-operative treatment | | | | |
| R + TMZ = "Stupp" | 5/15 | 34% | 41/55 | 75% |
| Radiotherapy only | 2/15 | 13% | 3/55 | 5% |
| Chemotherapy only | 4/15 | 27% | 2/55 | 4% |
| No treatment | 2/15 | 13% | 4/55 | 7% |
| NA | 2/15 | 13% | 5/55 | 9% |
| Overall survival (years) | | | | |
| Median | Not reached | | 1.25 | |
| Range | [0.1-6.6] | | [0.1-3.8] | |

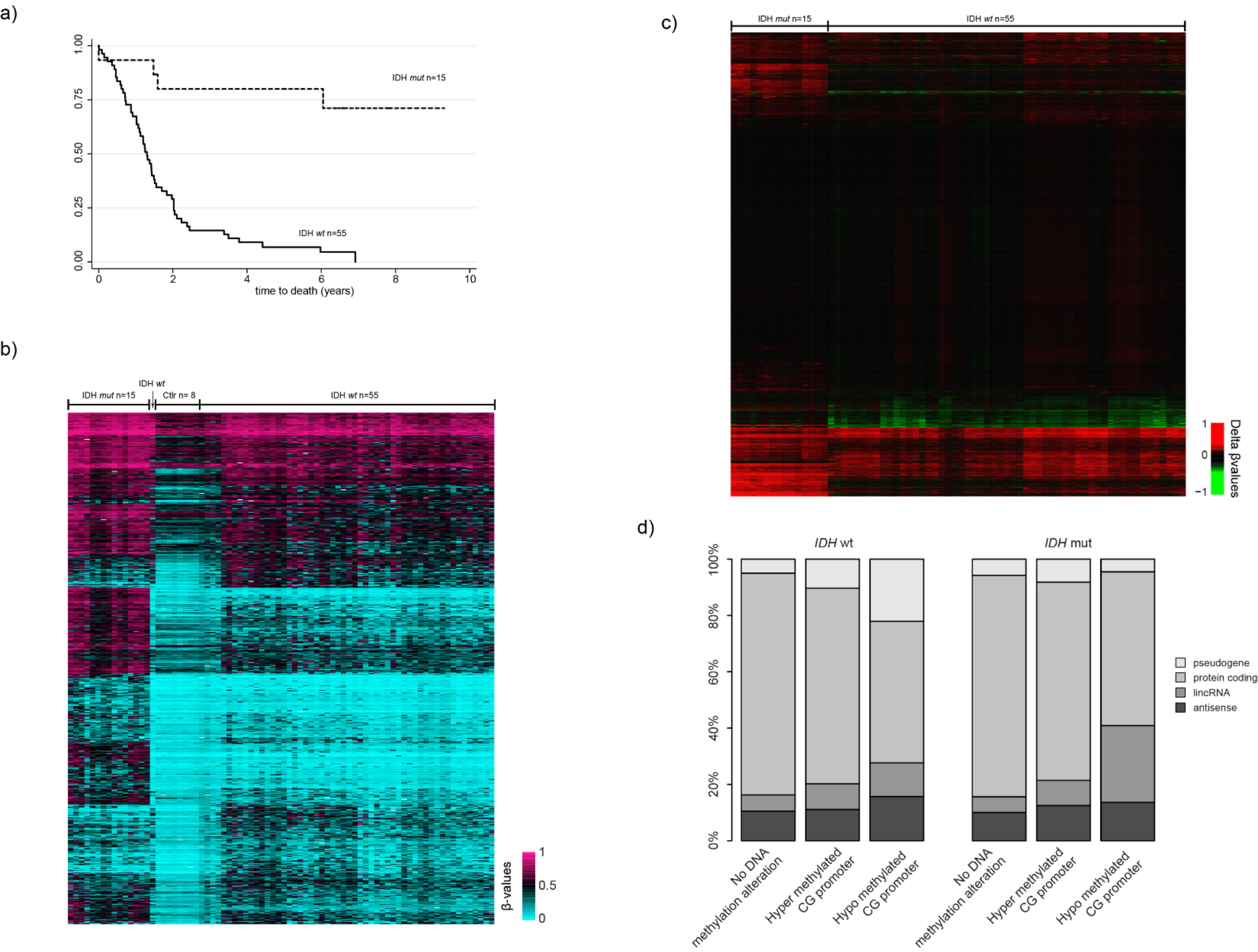


**Figure S1: DNA methylation alteration in IDHwt and IDHmut glioma samples**

**a)** Kaplan–Meier survival curves for patients with IDHwt (n=55) and IDHmut (n=15) glioma confirmed that the IDH status is a significant prognostic marker: 2-year survival rate of 27% in patients with IDHwt (median survival of 1.2 years/14.4 months) and 83% in patients with IDHmut glioma (median survival not reached). **b)** Visualization of the DNA methylation level (β-values) at the 86157 probes (row) located in the 11795 CGI/promoters analyzed in this study in IDHmut, controls and IDHwt glioma samples (column) identified a G-CIMP profile in IDHmut glioma samples. **c)** Differential DNA methylation level vs controls (delta of the means of β-values) of the 11795 CGIs (row) analyzed in this study in IDHmut and IDHwt glioma samples (columns). **d)** Classification of the genes associated with hyper-, hypo-methylated or unaffected CGI/promoters, respectively, in IDHwt (upper panel) and IDHmut (lower panel) glioma samples.

a)

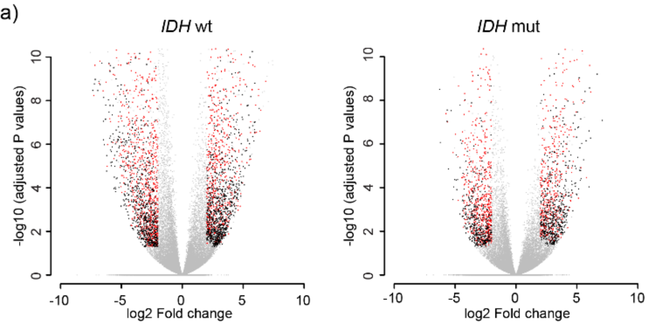


| | IDH wt Up fdr<0.05 & log2(FC)>2 | IDH wt Down fdr<0.05 & log2(FC)< -2 | IDH mut Up fdr<0.05 & log2(FC)>2 | IDH mut Down fdr<0.05 & log2(FC)< -2 |
|---|---|---|---|---|
| all Genes (44267) | 1865 / 4.2% | 1865 / 4.2% | 1085 / 2.5% | 1114 / 2.5% |
| **no CGI/promoter Genes (n=25850)** | **845 / 3.3%** | **748 / 2.9%** | **511 / 2%** | **386 / 1.5%** |
| CGI(s)/promoter Genes (n=18417) | 1020 / 5.5% | 1117 / 6.1% | 574 / 3.1% | 728 / 4% |
| **CGI/promoter Genes (n=14714)** | **781 / 5.3%** | **889 / 6%** | **460 / 3.1%** | **564 / 3.8%** |

b)

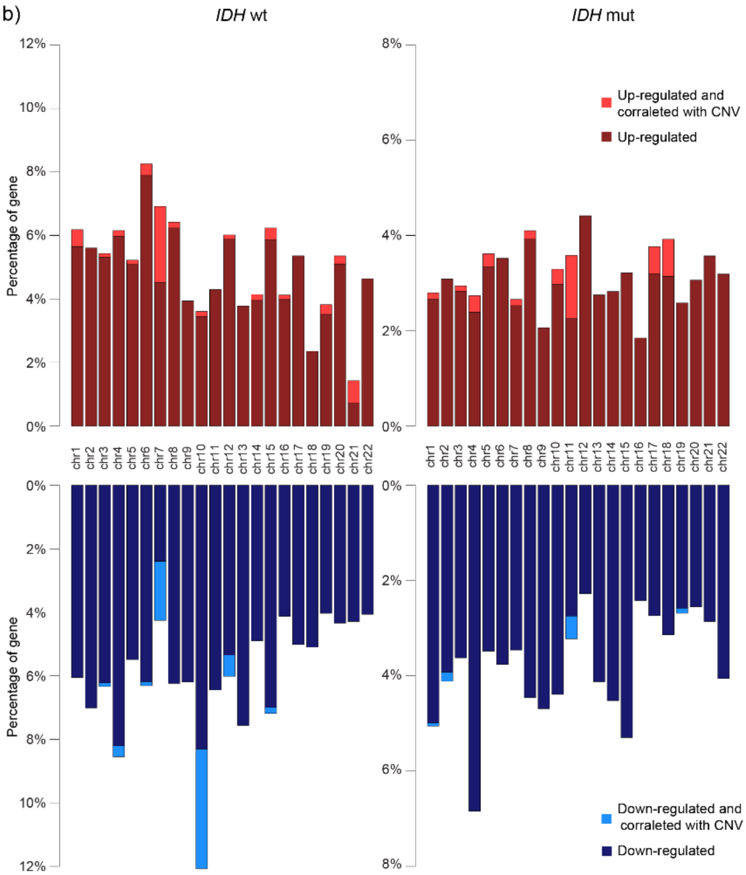


c)

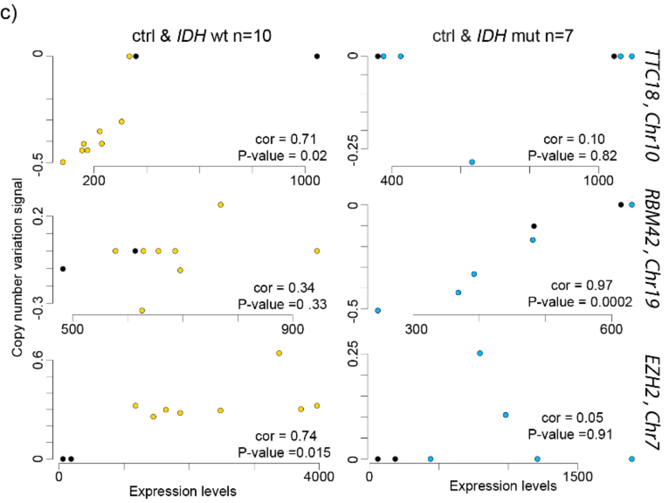


**Figure S2: Transcriptional alterations in IDHwt and IDHmut glioma samples.**

**a)** Volcano plot analysis with DESeq2 of the differential expression of all annotated genes (n= 44267) in IDHwt (left) and IDHmut (right) glioma samples compared with healthy brain controls. Genes that were significantly up- or down-regulated are symbolized by a red dot for CGI/promoter-associated genes, and by a black dot for the others. The numerical values are given in the table. **b)** Distribution per chromosome of transcriptionally up- (upper panels) and down-regulated (lower panels) CGI/promoter-associated genes in IDHwt and IDHmut glioma samples. The relative proportion of gene in which transcription alteration correlated with CNV is indicated. **c)** Details of the correlation analyses between CNV and expression for the *TTC18*, *EZH2* and *RBM42* genes in IDHwt (yellow dots in the left panels) and IDHmut (blue dots in the right panels) glioma samples. Black dots indicate the expression value in healthy brain. *TTC18* downregulation and *EZH2* overexpression correlated with CNV in IDHwt glioma samples. *RBM42* downregulation correlated with CNV in IDHmut glioma samples.

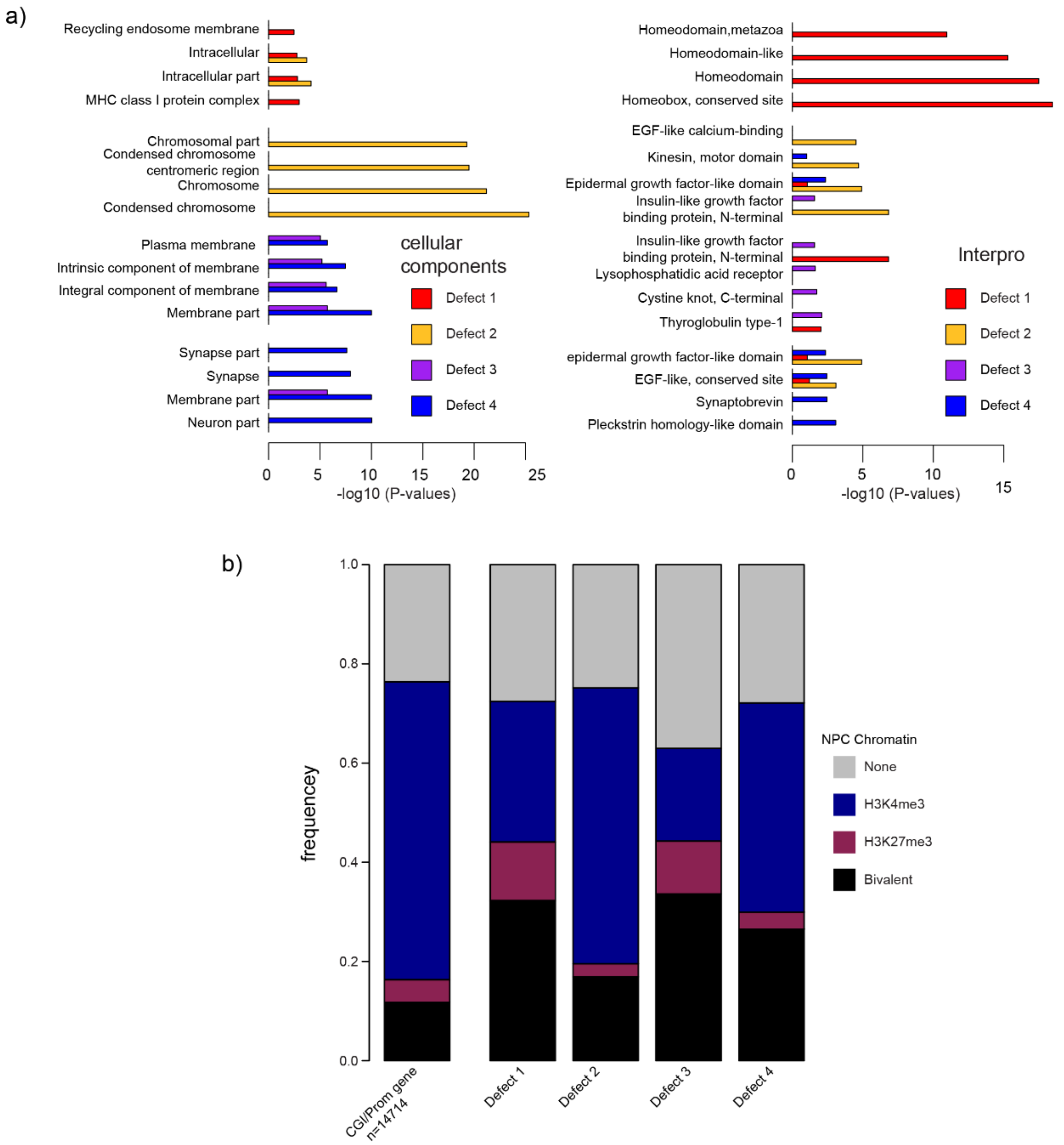


**Figure S3: Gene ontology**

**a**) Gene ontology terms enriched in genes with defect 1 to 4 identified in IDHwt glioma samples. For each category, the four highest terms are shown. **b**) Distribution of genes with defect 1 to 4 according to their chromatin signature in human NPC (none: gray; bivalent: black; H3K4me3-only: blue; H3K27me3-only: red). As reference, the distribution of the 14714 genes analyzed in this study according to their chromatin signatures in human ES cells is shown in the left panel.

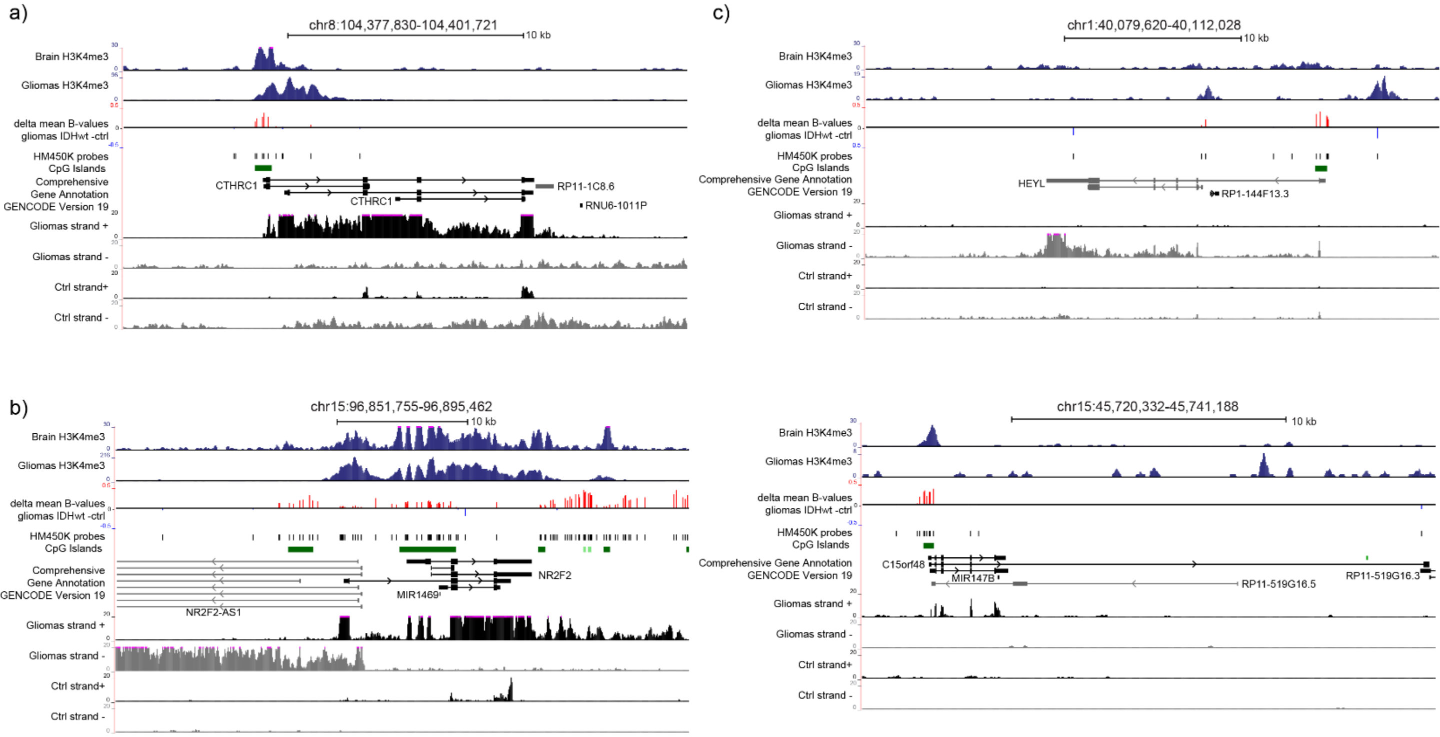


**Figure S4: Expression of genes with a methylated CGI/promoter**

Genome browser view showing H3K4me3 enrichment, differential DNA methylation, and oriented RNA-seq signal at **a**) *CTHRC1* (representative example of genes in which transcription initiates from an H3K4me3-marked TSS embedded in a methylated CGI/promoter, **b**) *NR2F2* (example of genes in which transcription initiates from an alternative TSS, and **c**) *HEYL* and *C15orf44*, where transcription initiates from a methylated CGI/promoter.

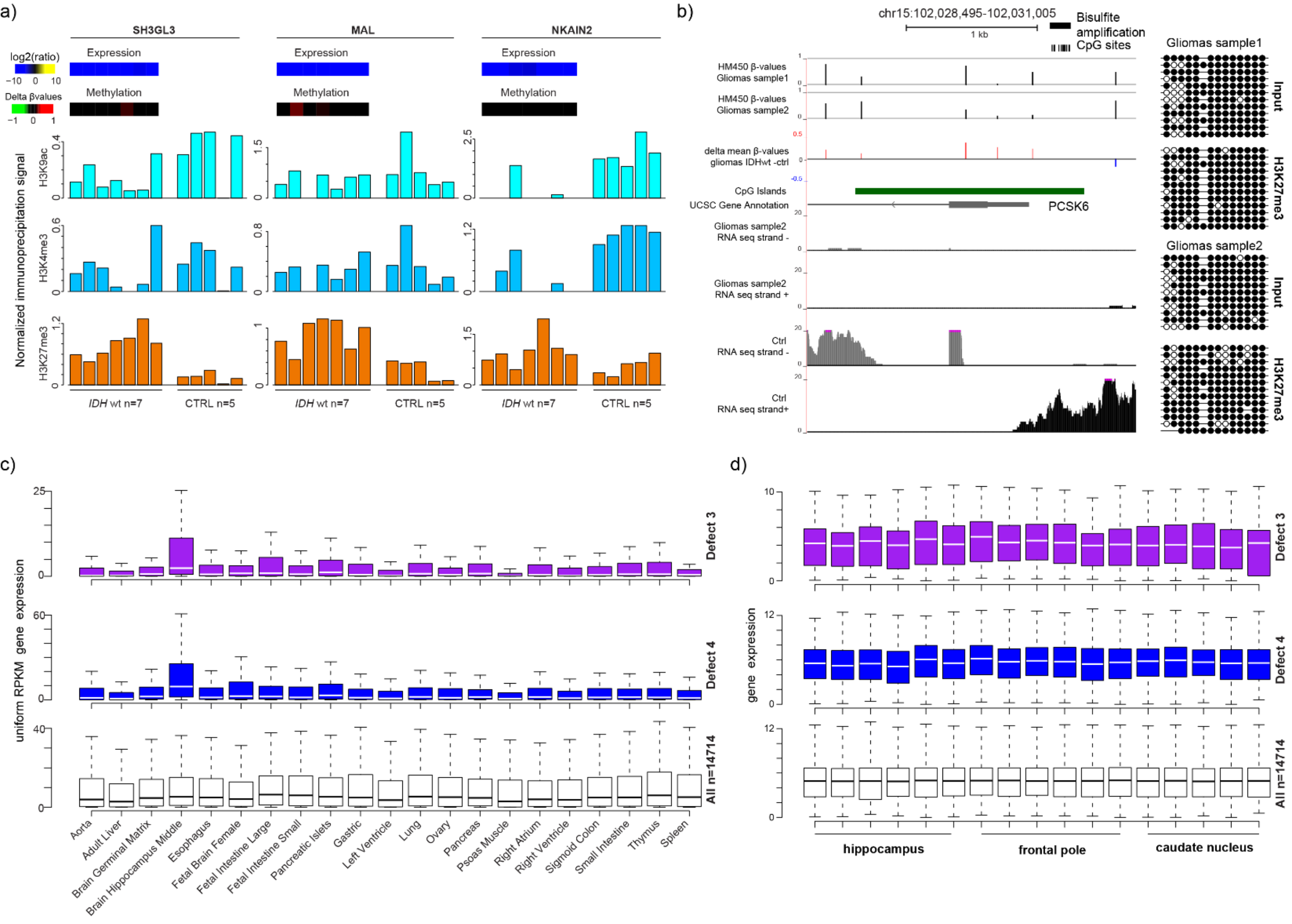


**Figure S5: Gene repression is associated with H3K27me3 gain and affect mainly genes with brain-specific expression profile**

**a)** ChIP analysis of H3K9ac, H3K4me3 and H3K27me3 at selected genes in IDHwt glioma (n=7) and control (n=5) samples. The precipitation level was normalized to that obtained at the *TBP* promoter (for H3K4me3 and H3K9ac) and at the *SP6* promoter (for H3K27me3). **b)** Bisulfite-based sequencing data for the *PCSK6* CGI/promoter using input and H3K27me3-immunoprocipated ChIP fractions showed that DNA methylation and H3K27me3 can coexist in glioma cells. Each horizontal row of circles represents the CpG dinucleotides on an individual chromosome. Solid circles, methylated CpG dinucleotides; open circles, unmethylated CpG dinucleotides. The relative position of the bisulfite amplicon is showed on the *PCSK6* locus browser view (right panel). **c)** Median expression of genes with defect 3 (purple column), 4 (blue column) or all CGI/promoter genes (white column) in 21 tissues (publicly available normalized RNA-seq data). Genes with defect 4 are strongly expressed specifically in adult hippocampus **d**) The expression level in hippocampus is representative of the expression level in other brain parts, as shown for the frontal pole and caudal nucleus.

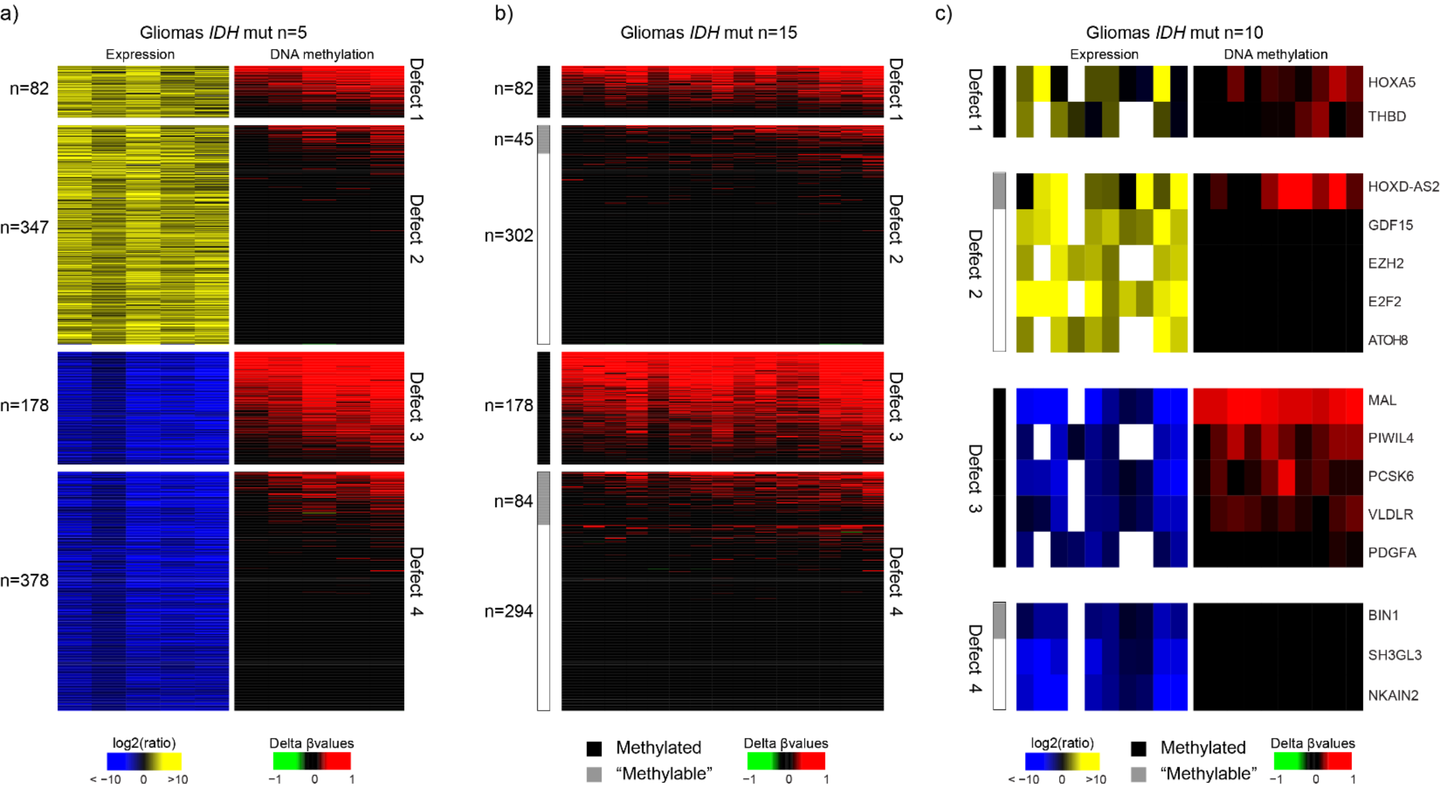


**Figure S6: Four classes of expression defects in IDHmut glioma samples**

**a)** Integrative analysis of gene expression and methylation assessed in 5 IDHmut glioma samples identified four main defect classes. **a)** Differential DNA methylation analysis of all the IDHmut glioma samples (n=15) classified according the defect classes defined in a). The methylated and methylable status of the genes is indicated in the left column. **c)** Integrative analysis of the differential expression and methylation (vs control) for selected genes from each class defect in 10 IDHmut glioma samples.

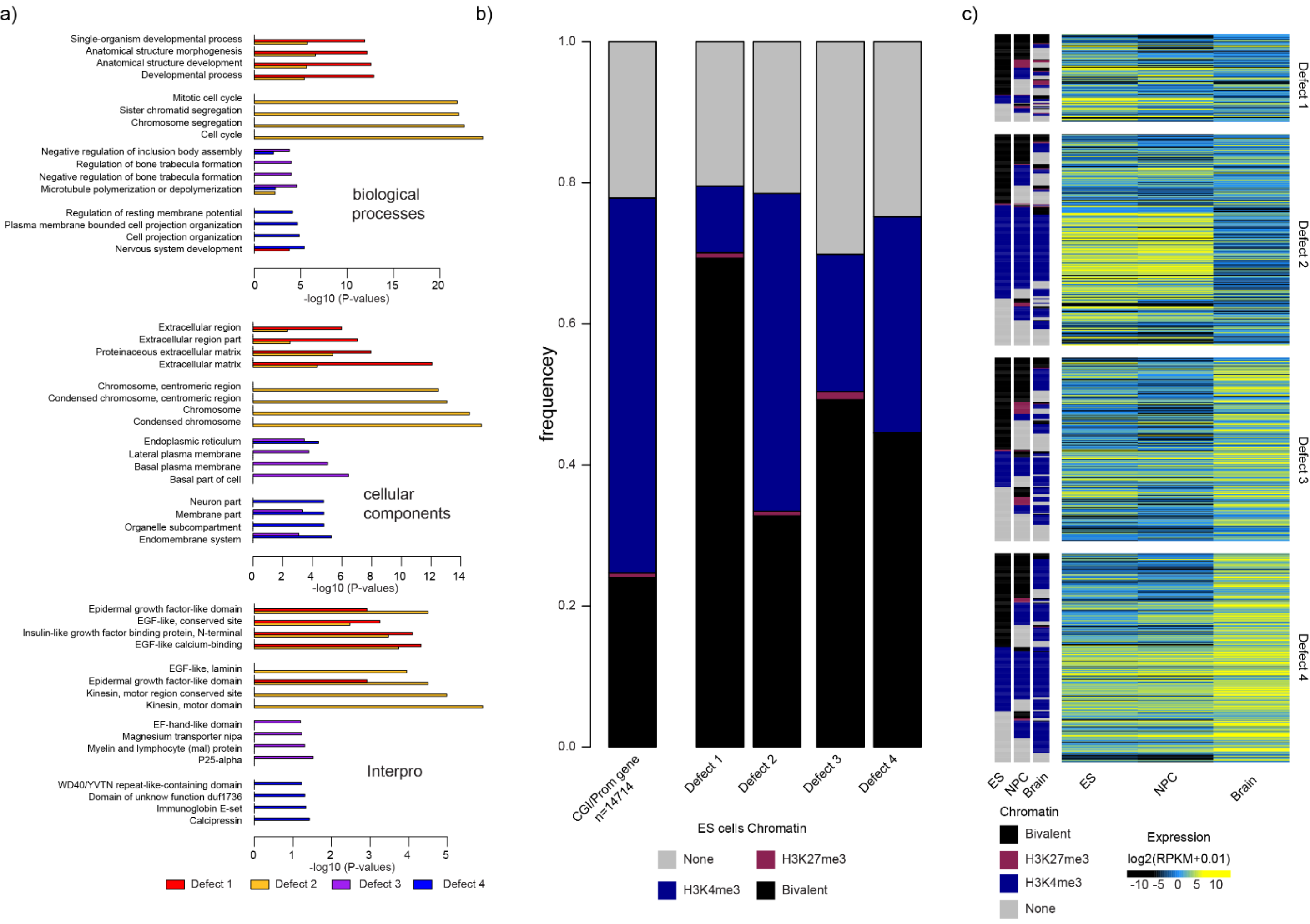


**Figure S7: Genes with bivalent chromatin signature in ES cells are more prone to be deregulated in IDHmut glioma.**

**a)** Gene ontology terms enriched in genes with defect 1 to 4 identified in IDHmut glioma samples. For each category, the four highest terms are shown. **b)** Distribution of genes with defects 1 to 4 according to their chromatin signatures in human ES cells (none: gray; bivalent: black; H3K4me3-only: blue; H3K27me3-only: red). As a reference, the distribution according to their chromatin signatures in human ES cells of all the 14714 genes analyzed in this study is shown in the left panel. **c)** Expression level and associated chromatin signatures of genes with defects 1 to 4 in human ES cells, neural progenitors cells (NPC) and healthy brains.
